# Allosteric Inhibition of PTP1B by a Nonpolar Terpenoid

**DOI:** 10.1101/2022.05.18.492571

**Authors:** Anika J. Friedman, Evan T. Liechty, Levi Kramer, Ankur Sarkar, Jerome M. Fox, Michael R. Shirts

## Abstract

Protein tyrosine phosphatases (PTPs) are promising drug targets for treating a wide range of diseases such as diabetes, cancer, and neurological disorders, but their conserved active sites have complicated the design of selective therapeutics. This study examines the allosteric inhibition of PTP1B by amorphadiene (AD), a terpenoid hydrocarbon that is an unusually selective inhibitor. Molecular dynamics (MD) simulations carried out in this study suggest that AD can stably sample multiple neighboring sites on the allosterically influential C-terminus of the catalytic domain. Binding to these sites requires a disordered *α*7 helix, which stabilizes the PTP1B-AD complex and may contribute to the selectivity of AD for PTP1B over TCPTP. Intriguingly, the binding mode of AD differs from that of the most well-studied allosteric inhibitor of PTP1B. Indeed, biophysical measurements and MD simulations indicate that the two molecules can bind simultaneously. Upon binding, both inhibitors destabilize the *α*7 helix and disrupt hydrogen bonds that facilitate closure of the catalytically essential WPD loop. These findings indicate that AD is a promising scaffold for building allosteric inhibitors of PTP1B and illustrate, more broadly, how unfunctionalized terpenoids can engage in specific interactions with protein surfaces.

## Introduction

Protein tyrosine phosphatases (PTPs) are an influential class of regulatory enzymes that have long eluded drug design; they are often referred to as “undruggable”, largely as a result of the low bioavalability and poor selectivity of inhibitors. These enzymes regulate cellular growth, motility, and oncogenic transformation, and contribute to a broad set of complex physiological processes (e.g., memory, inflammation, metabolism, and autoimmunity).^1–8^ Classical PTPs catalyze the hydrolytic dephosphorylation of tyrosine residues with four conserved active site loops: (i) the P-loop (C(X)5R(S/T)), where an arginine facilitates substrate binding and transition state stabilization and a cysteine enables nucleophilic attack of the phosphate ester, (ii) the WPD loop (10-12 residues), which has the general acid catalyst required for hydrolysis, (iii) the Q-loop, where a glutamine positions a water for nucleophilic attack of the phosphocysteine intermediate, and (iv) the substrate binding loop, which selects for phosphorylated tyrosine residues.^9^ The conserved active site of PTPs has hindered the design of selective therapeutics.

The catalytic domains of PTPs include allosteric networks that communicate between the active site and less conserved regions.^10,11^ Protein tyrosine phosphatase 1B (PTP1B) provides an illustrative example.^12–14^ Over the years, a myriad of biophysical analyses of this enzyme have yielded two particularly important findings: (i) the closure of its WPD loop enables dephosphorylation of the phosphocysteine intermediate, a rate-limiting step in catalysis, and (ii) this motion is regulated by a network of hydrogen bonds (h-bonds) that extends to the C-terminal *α*7 helix on the catalytic domain.^10,15–20^ Interactions between the *α*3, *α*6, and *α*7 helices—referred to in this paper as the helical triad—affect the intermediate timescale dynamics of WPD loop motion. At conformational extremes, an ordered *α*7 helix stabilizes a closed WPD loop, and a disordered helix stabilizes an open WPD loop. The removal of the *α*7 helix reduces catalytic activity by 40-60%.^21,22^

Allosteric inhibitors that bind to poorly conserved sites on PTP1B are promising starting points for building selective therapeutics. An early screen identified benzobromarone derivatives that bind outside of the active site.^23^ These inhibitors displace the *α*7 helix, restrict rotation of the *α*3 helix, and prevent the formation of h-bonds that stabilize a closed WPD loop.^22^ Unfortunately, these molecules have not been translated into approved drugs. Nuclear magnetic resonance (NMR) analyses and multi-temperature crystallography have uncovered other allosterically influential regions—most notably, the disordered C-terminus that extends from the catalytic domain to the endoplasmic reticulum, the “197 site”, which sits between the *α*3 helix and a *β*-sheet, and the “L16” site (residues 237-243), located beneath the *α*6-α7 junction.^24,25^ The design of inhibitors that bind to these regions, however, remains challenging and to date, no inhibitors of PTP1B have entered phase III clinical trials.^5,26–32^

Motivated by the paucity of well-characterized allosteric inhibitors of PTPs, we used an engineered microbial system to search for terpenoids that inhibit PTP1B.^33^ We reasoned that nonpolar terpenoids, if inhibitory, would bind outside of the positively charged active site. Indeed, kinetic analyses and X-ray crystallography showed that amorphadiene (AD) can inhibit PTP1B by binding to a hydrophobic pocket formed by a reorganization of its *α*7 helix. AD is a surprisingly selective and potent inhibitor for a small, “greasy” molecule (Figure 1A); its IC_50_ is approximately 50 *μ*M, and it inhibits PTP1B 5- to 6-fold more potently than TCPTP, which shares 69% sequence identity.^34^ AD also appears to engage in loose, conformationally flexible binding, a behavior evidenced experimentally by ill-defined regions of electron density around its crystallographic binding site.^33^

**Figure 1:**
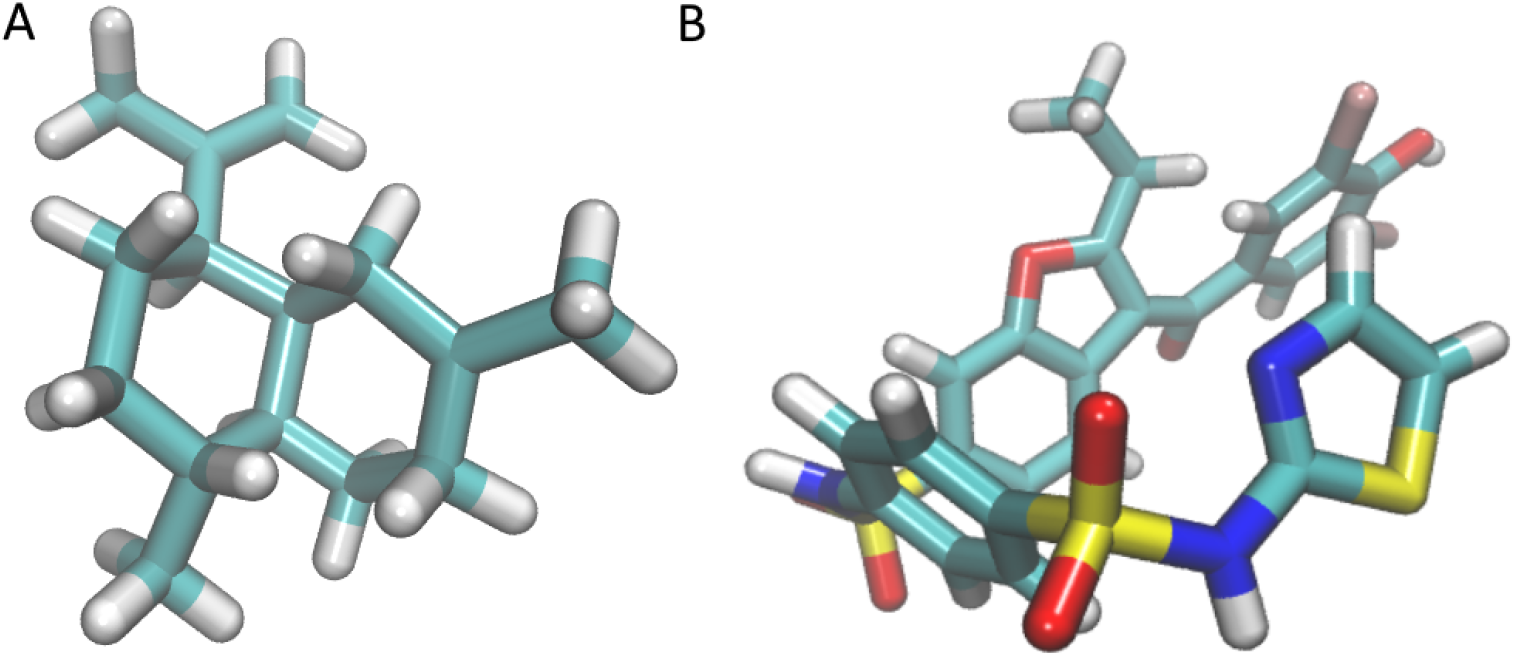
The chemical structure of AD (A) and BBR (B) are distinct. AD is significantly smaller in size and lacks the h-bond donors and acceptors that allow BBR to form stabilizing h-bonds. Given the structure of AD, it is surprising that AD exhibits similar selective binding to PTP1B over TCPTP and inhibits PTP1B with an IC50 only ~ 6x greater than BBR^23,33^

This study combines molecular dynamics (MD) simulations, detailed kinetic measurements, and binding analysis to study the mechanism by which AD inhibits PTP1B. It focuses on three important questions: (i) How does AD interact with the *α*7 helix to form a stable complex? (ii) How does the PTP1B-AD complex disrupt enzyme activity? (iii) How do mechanisms of inhibition differ between AD and previously characterized benzobromarone derivatives? Answers to these questions could reveal new varieties of allosterically influential interactions with PTP1B, illustrate how unfunctionalized terpenoids can engage in specific interactions with protein surfaces, and inform the design of new inhibitory compounds.

## Materials and Methods

### Materials

We used chemically competent NEB Turbo *E. coli* cells for cloning and BL2(DE3) *E. coli* cells to express PTP1B (New England Biolabs). We purchased 3-(3,5-Dibromo-4-hydroxybenzoyl)-2-ethyl-benzofuran-6-sulfon-icacid-(4-(thiazol-2-ylsulfamyl)-phenyl)-amide (BBR) and 2-[(Carboxy-carbonyl)amino]-4,5,6,7-tetrahydrothieno[2,3-c]pyridine-3-carboxylic acid hydrochloride (TCS 401) from Cayman Chemical (Ann Arbor, Michigan) and Ertiprotafib from Med-Koo Biosciences. We purchased HEPES Buffer (1 M 4-(2-hydroxyethyl)-1-piperazineethanesulfonic acid, pH 7.3) from Fisher, biochemical reagents (e.g., DNA polymerase) from New England Biolabs, and DNA primers from Integrated DNA Technologies. We isolated AD from microbial cell cultures as described previously.^33^

### Cloning and Molecular Biology

We constructed mutants of PTP1B with Gibson assembly. We designed all primers to have 60° C annealing temperatures and full complementary to facilitate plasmid assembly. We introduced point mutations near the middle of each primer to ensure proper annealing. We ligated all DNA segments at 50° C for one hour, and confirmed the presence of targeted mutations using Sanger Sequencing (QuintaraBio). Table S1 lists all primers.

### Protein Expression and Purification

We overexpressed mutant and wild-type forms of PTP1B (residues 1–321) on pET16b plasmids, where the PTP1B gene was fused to a C-terminal 6x polyhistidine tag. We transformed E. coli BL21(DE3) cells with each pET16b vector and grew the transformed cells in 1-L cultures to an *OD*_600_ of 0.5-0.8 (37° C, 225 RPM), induced them with 500 *μ*M IPTG, and grew them at 22°C for 18 hours. We lysed the cells with lysis buffer (20 mM Tris HCl, 50 mM NaCl, 1% Triton X-100, pH 7.5) and purified PTP1B with nickel affinity and anion exchange chromatography (HisTrap HP and HiPrep Q HP, respectively; GE Healthcare). We used 10,000 MW cutoff spin columns for each buffer exchange (Satorius). We stored the final protein in HEPES buffer (50 mM, pH 7.5, 0.5 mM TCEP) in 20% glycerol at 80° C.

### Analysis of Binding Affinity

We examined the binding of inhibitors to PTP1B by measuring binding-induced changes in tryptophan fluorescence. In brief, we measured the fluorescence (280_ex_/370_em_) of 5 *μ*M PTP1B in the presence of 0-500 *μ*M of BBR (50 mM HEPES, 8% DMSO, pH=7.3) under three conditions: (i) BBR alone, (ii) BBR with 115 *μ*M AD, and (iii) BBR with 30 *μ*M TCS401. These concentrations of AD and TCS401 produce similar levels of inhibition (i.e., ~50%; Fig. S1). To assemble binding isotherms, we fit fluorescence data to Δ*F* = (Δ*F_max_*\*L)/(*K_d_*+L), where Δ*F* is the change in tryptophan fluorescence caused by BBR, L is the concentration of BBR (in *μ*M), Δ*F*_max_ is the maximum change in fluorescence, and *K_d_* is the dissociation constant. Table S3 reports raw fluorescence data, calculated values of Δ*F*, and final fit parameters (i.e., Δ*F_max_* and *K_d_*).

### Differential Scanning Fluorimetry (DSF)

We used DSF to examine the influence of several inhibitors (AD, BBR, Ertiprotafib, and TCS401) on the melting temperature of PTP1B. We dissolved each inhibitor in 100% DMSO at 50X the desired concentration (0–300 *μ*M) and pre-incubated 1 *μ*L of this solution with 49 *μ*L of protein solution [(2 *μ*M PTP1B, 50 mM HEPES (pH 7.3), 5X SYPRO Orange dye (Life Technologies, Eugene, OR)] for at least ten minutes; here, we ensured that the maximum inhibitor concentration reached at least 3× the IC_50_. We used a StepOnePlus RT-PCR instrument (Life Technologies, Eugene, OR) to perform a melting curve analysis with detection settings for the Rox reporter (580*_ex_*/621*_em_* nm) and the following temperature regime: hold at 25° C (2 min), ramp to 95°C at 1° C/min, and hold at 95° C (2 min). We exported final temperature, normalized fluorescence, and first-derivative data for the melt region (Figure S2), and estimated melting temperatures (T*_m_*) by calculating the local minima of the negative first-derivative data (Figure 2C).^35^

**Figure 2:**
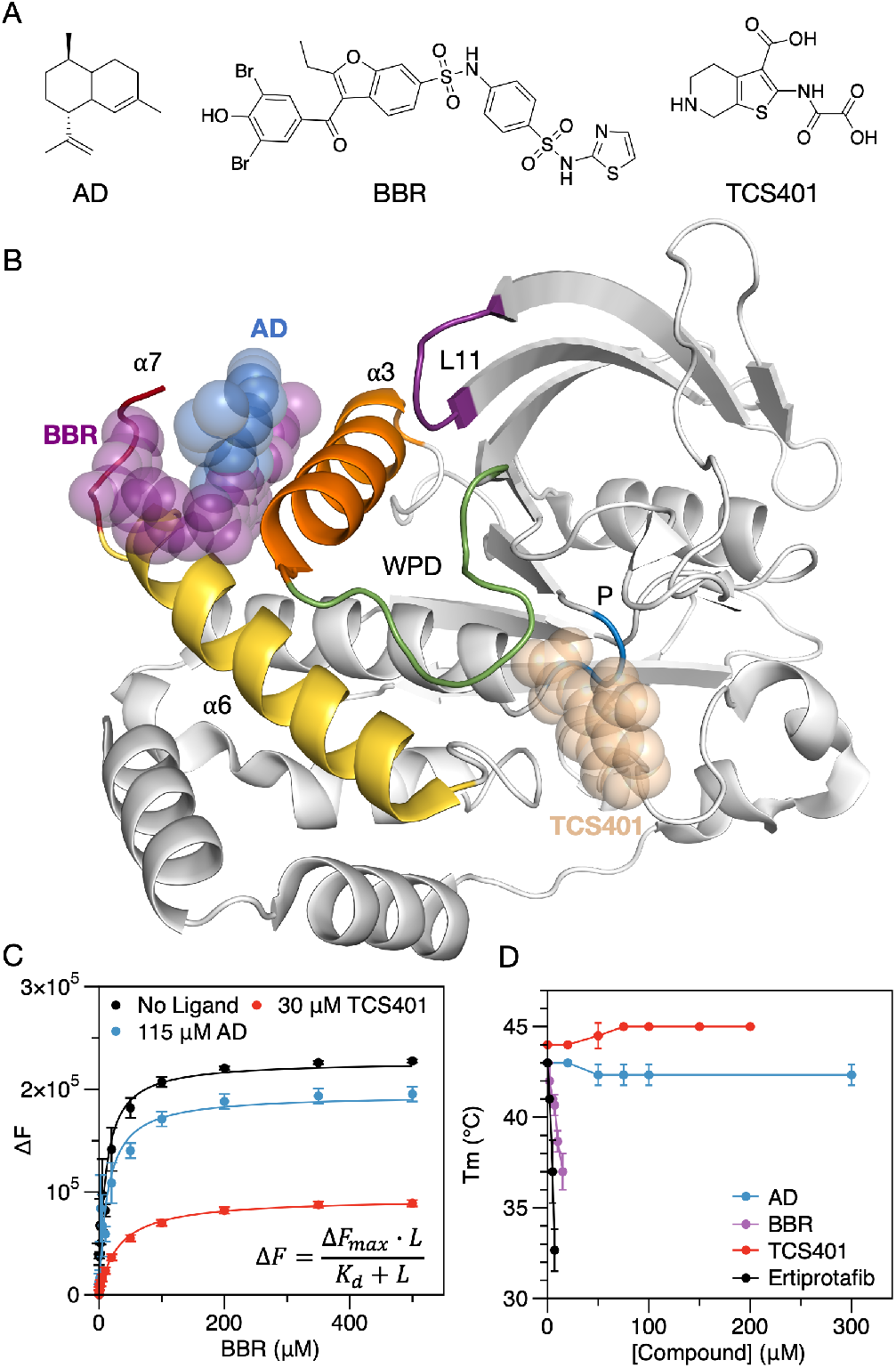
(A) Structures of amorphadiene (AD) and well-studied allosteric (BBR) and competitive (TCS401) inhibitors. (B) An X-ray crystal structure of PTP1B bound to AD (PDB entry 6W30) with the binding sites for BBR and TCS401 overlaid for reference. We aligned structures of the PTP1B-AD, PTP1B-BBR, and PTP1B-TCS401 complexes (PDB entries 6W30, 1T4J, and 5K9W) by using the “align” function from PyMol. AD and BBR bind to the allosteric site, which includes residues from the *α*3, *α*6, and *α*7 helices; TCS401 binds to the active site, which is flanked by the WPD and P-loops. (C) Fluorescence-based binding isotherms for BBR measured in the presence and absence of either AD or TCS401. We ensured similar levels of binding by AD and TCS401 by using concentrations that produced similar levels of inhibition (~50%; Figure S1). Binding parameters (± SE) for Δ*F* = (Δ*F_max_*\*L)/(*K_d_*+L): *K_d_* = 10.1 ± 2.7 *μ*M and Δ*F_max_* = 227000 ± 13000 (BBR alone); *K_d_* = 13.1 ± 3.8 *μ*M and Δ*F_max_* = 195000 ± 12000 (BBR with AD); *K_d_* = 31.0 ± 2.8 *μ*M and Δ*F_max_* = 94000 ± 2000 (BBR with TCS401). The insensitivity of the BBR binding isotherm to the presence of AD suggests that the two inhibitors can bind simultaneously. Error bars denote standard error for *n* = 3 technical replicates. (D) Melting temperatures determined with differential scanning fluorimetry indicate that BBR and Ertiprotafib destabilize PTP1B, while AD and TCS401 do not. Error bars denote standard deviation for *n* = 3 technical replicates.

### Molecular Dynamics (MD) Simulations

We prepared PTP1B for MD simulations by starting with three X-ray crystal structures: (i) apo PTP1B with an ordered *α*7 helix and a closed WPD loop (PDB code: 1SUG), (ii) PTP1B in complex with BBR (PDB code: 1T4J), and (iii) PTP1B in complex with AD (PDB code: 6W30).^23,33,36^ Both protein-ligand complexes had an open WPD loop and a disordered *α*7 helix, which prevented resolution with X-ray crystallography. For each structure, we removed crystallized waters, glycerol, and Mg^2+^, adjusted the protonation state to a pH of 7 using the H++ web-server, added Na^+^ ions to neutralize the net charge, and hydrated the protein with a TIP3P water box, maintaining a minimum distance of 10 Å between the protein or ligand and the periodic boundary.

We carried out MD simulations with GROMACS 2020.4^37^ using Bridges-2 from the Pittsburgh Supercomputing Center. In all simulations, we modeled PTP1B with the AMBER *ff99sb-ildn* force field and parameterized AD and BBR with the Open Force Field v.1.3.0 “Parsley”).^38^ All analysis scripts and input parameters can be found in the repository at https://github.com/shirtsgroup/PTP1B. Ligand parameterization scripts can be found in repository folder “Ligand Parameters”. We carried out an energy minimization to 100 kJ/mol/nm force tolerance, and equilibrated the protein in the NVT ensemble at 300 K for 100 ps, followed by equilibration to the NPT ensemble at 300 K and 1 atm for 100 ps. All simulations used the velocity rescaling thermostat^39^ and Beredensen weak-coupling barostat. Further configuration details for the simulations appear in the repository folder “data/mdp”. We ran all MD simulations for 300 ns (unrestrained NPT) and visualized MD trajectories with Visual Molecular Dynamics 1.9.3 (VMD).^40^

X-ray crystal structures of PTP1B bound to BBR or AD lack the *α*7 helix, which becomes partially disordered when PTP1B binds to these inhibitors. This conformational disorder prevents resolution with X-ray crystallography. Previous mutational analyses suggest that the disordered *α*7 helix mediates interactions between both inhibitors and PTP1B.^22^ Accordingly, for a subset of simulations, we reconstructed the *α*7 helix (i.e., residues 287–299) using Modeller 10.1 and a reference structure with the helix ordered and intact (PDB 1SUG). We reconstructed missing residues into an ordered helix using homology modelling to fit the structure of the known sequence of C-terminal residues (residues 280–299; code can be found in the “build_a7” folder of the repository). We then generated an ensemble of disordered helical conformations using restrained heating and allosteric ligand binding. We applied positional restrains (1000 kJ/mol/nm^2^ on all atoms) to all protein residues outside of the *α*7 helix (residues 1–280), heated the system gradually from 400 K to 500 K over 300 ns, and selected three disordered conformations from the final 50 ns of simulations, where the helix was completely disordered (i.e., the Defined Secondary Structure Prediction (DSSP) algorithm labeled 0% of residues *α* helical). We supplemented these disordered conformations with a fourth, which we selected at random from the final 25 ns (in which the helix was fully disordered) of our simulation of the PTP1B-BBR complex initialized with an ordered *α*7 helix and an open WPD loop.

We chose a variety of starting configurations to probe the effects of the introduction of the ligand to the structure of PTP1B. We initialized simulations of apo and AD-bound PTP1B with the WPD loop in open (derived from 6W30; WPD_open_) and closed (derived from 1SUG; WPD_closed_) conformations and with the *α*7 helix ordered, disordered, and absent (Table S2). The WPD_closed_ conformation allowed us to determine the conformational changes induced by the introduction of the ligand to the binding site; however, the timescales of these simulations were potentially insufficient for PTP1B-ligand complex to reach a stable final equilibrium. The WPD_open_ conformation was used to reduce the conformational changes required for the ligand-bound structure to equilibrate and achieved stable equilibrium on the timescale of the simulations. For BBR, we ran the same simulations used for AD, excluding those lacking the *α*7 helix. Previous studies have elucidated the importance of this helix in BBR binding, so we included only *α*7-containing structures in our analysis of the PTP1B-BBR complex.

### Analysis of MD Trajectories

Before completing analysis on our MD trajectories in detail, we carried out two important processing steps: (i) removal of correlated trajectory frames (ii) removal of unequilibrated trajectory frames and determination of convergence. Correlated trajectory frames were removed with ruptures 1.1.6^41^ and unequilibrated trajectory frames were removed based on the root-mean-square deviation (RMSD) of backbone atoms, relative to the starting structure for the production simulation (Further details in sections S1.1 and S1.2).

Our MD trajectories suggested that AD can bind to several different sites on PTP1B. We classified these binding sites as follows (also pictured in Figure 3A–B):

1. At loc1, the crystallographic binding location, AD engages in simultaneous interactions with the *α*3 and *α*7 helices without interacting with the *α*4 and *α*5 helices or the N-terminal region of the *α*6 helix (residues 264–270);
2. At loc2, AD interacts with the *α*6 and *α*7 helices without interacting with the *α*4 or *α*5 helices and engages in no more than one interaction with the *α*3 helix;
3. At loc3, AD interacts with the *α*4 and *α*6 helices without interacting with the *α*3 or *α*7 helices;
4. At loc4, AD interacts with the *α*3, *α*4, and *α*6 helices without interacting with the *α*5 or *α*7 helices.

**Figure 3:**
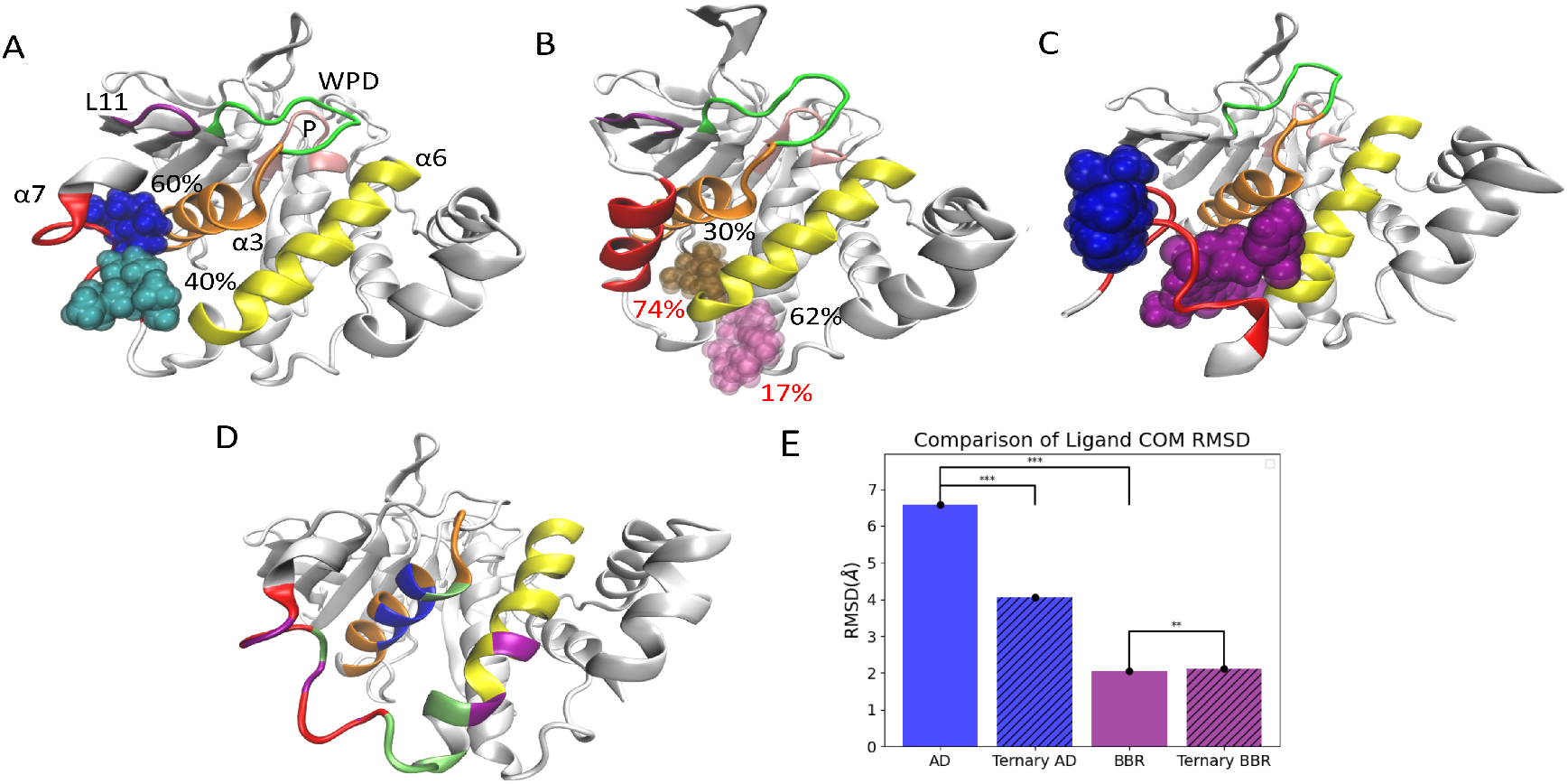
AD is capable of occupying a diverse set of binding conformations. (A) In MD simulations initialized with a disordered *α*7 helix, AD samples two adjacent sites with near-equal frequency: the crystallographic site (loc1; blue) and a neighboring site (loc2; turquoise). The percent occupancy of both sites are shown in the figure (details on determination of occupancy percentages in section S1.6). (B) When the *α*7 helix is initialized with an ordered (black percentages) conformation or absent (red percentages), AD moves to two new sites: loc3 (pink) and loc4 (brown). (C) In MD simulations initialized with AD and BBR at their crystallographic binding sites, AD moves to the outside of the *α*7 helix and remains at this location (blue) for the entire duration of the 1 *μ*s trajectory. In A–C, the protein and ligand represent centroid structures from the corresponding MD trajectories. (D) During MD trajectories, AD (bound to loc1 and loc2) and BBR interact with the same core set of residues (green) and several residues specific to AD (blue) or BBR (purple). (E) A comparison of RMSDs for the COM of AD and BBR in different complexes. AD exhibits significantly higher fluctuations than BBR in complex with PTP1B, a result of frequent oscillations by AD between adjacent binding locations. In the ternary complex, AD exhibits significantly reduced COM motion, as it no longer oscillates between binding locations. BBR experiences a 3% increase in COM RMSD in the ternary complex; this change is statistically significant but very small, relative to differences in the COM RMSDs of BBR and AD, or AD in the presence and absence of BBR.

We considered the ligand to be unbound when it had no interactions with the *α*3–α7 helices. Any frames in which the ligand was bound but not to one of our defined binding sites were classified as “other bound” states. These were unstable and not conserved between trajectories and were thus not otherwise classified. In the above, we define interactions as a distance of <5 Å between heavy atoms in the ligand and protein residues, a relatively generous distance.

The WPD loop of PTP1B can adopt an open or closed conformation. We classified its position by the distance between the *α*-carbons of D181 and C215 (i.e., the catalytic acid and the nucleophile, respectively),^17,42,43^ as measured using using the compute_distances function of MDTraj. Crystal structures with the WPD loop in closed and open states had D181–C215 distances of 8 and 15 Å, respectively, so we used a distance of 10 Å to differentiate between states: WPD_closed_ (< 10 Å) and WPD_open_ (≥ 10 Å). As confirmation, the combination of distances from all MD trajectories showed a bimodal distribution with a minimum at approximately 10 Å (Figure S3D).

We examined the helicity of the *α*7 helix in our MD trajectories by using the DSSP algorithm implemented in MDTraj 1.9.4.^44^ This algorithm characterizes the secondary structure of each residue based on the *ϕ* and *ψ* torsional angles. Importantly, DSSP can differentiate between different types of helices: *α* helix, 3_10_ helix, and *π* helix. This analysis allowed us to characterize the order, or lack thereof, of the *α*7 helix. In this paper, “α helicity” is specific to residues with an *α* helix conformation while “helicity” alone generalizes to include all listed helix types.

To further classify the structure of PTP1B throughout the simulations, we evaluated the RMSD of the backbone atoms and the root-mean-square-fluctuation (RMSF) of select protein regions, relative to a centroid structure. We defined the centroid structure by clustering each trajectory on the backbone atoms of the equilibrated trajectory using gmx_cluster and taking the centroid of the cluster consisting of all structures. For each trajectory, we evaluated the RMSD of the backbone atoms relative to both (i) the centroid structure for the trajectory and (ii) the centroid structure for the trajectory of the apo protein initialized with the WPD loop in the closed conformation (with an ordered *α*7 helix). The second analysis allowed us to search for structural changes in the the protein induced by inhibitor binding. For ligand bound trajectories the center-of-mass (COM) RMSD for the ligand was also computed using bootstrapping on the uncorrelated configurations to determine the mean and standard error of the ligand COM RMSD value (further details in section S1.3).

The catalytic domain of PTP1B has seven *α*-helices that play an important role in allosteric communication. We quantified inter-helical interactions and helix-ligand interactions between these influential helices as those with a residue-residue or residue-ligand distance of less than 5 Å. We defined inter-helical interactions disrupted by ligand binding as those that occur significantly less (*p* < 0.05) in the ligand bound vs. corresponding apo conformation. We calculated the p-value using Welch’s T-test for the fraction of the simulation time that the interaction was present for the ligand bound (AD or BBR) compared to apo trajectories. For this analysis ligand bound trajectories all maintain an open WPD loop and feture a disordered *α*7 helix with the ligand bound in its crystallographic pose (or in loc2 for AD).

We isolated allosterically influential h-bonds with several steps. (i) We used the Baker-Hubbard model implemented within `MDTraj` to identify h-bonds. This model uses a proton donor-acceptor distance of 2.5 Å and a donor-acceptor angle of less than 120° to classify h-bonds. (ii) We removed h-bonds formed in a majority of all trajectories, regardless of WPD loop conformation or the presence of an allosteric inhibitor, or formed between adjacent (within 3) residues and calculated the percent of the trajectory in which each of the remaining bonds appeared. (iii) For each h-bond, we determined the mean frequency formed for four groups: Apo WPD_open_, Apo WPD_closed_, AD bound, and BBR bound. (iv) We identified bonds that showed a statistically significant (*p* < 0.01) difference between the groups. (v) Using our statistical threshold, we selected bonds that appeared more in either apo WPD_open_ or apo WPD_closed_ (with a minimum appearance of 70% in their primary state) to define h-bonding networks in each of these conformations. Notably, no h-bonds appeared significantly more or less frequently (given the above selection criteria) with ligands bound than in the apo WPD_open_ state.

### Analysis of Influential Mutations

We used our MD trajectories to build a list of mutations likely to modify interactions between PTP1B and each allosteric inhibitor. To begin, we selected a subset of residues that (i) showed more interactions with AD than BBR, or vice versa, or that (ii) previous studies suggested would influence binding. Residues 192, 195, and 196 in the *α*3 helix preferentially interact with AD while residues 276, 279, 286, and 287 in the *α*6 and *α*7 helices preferentially interact with BBR. F196 was mutated alanine to reduce the size of the hydrophobic side chain and eliminate the possibility of the formation of *π*-stacking interactions with the ligand. L192 and L195 were mutated to alanine, phenylalanine, and asparagine to explore the effect of reducing the size of the hydrophobic side chain, introduction of a phenol ring, and introduction of a polar side chain to the hydrophobic cleft. The E276F mutation replaces the negatively charged side chain with a phenol ring to increase the hydrophobicity of the allosteric pocket and potentially provide an additional stabilizing pi-stacking position. S286A and K279M mutations were also intended to increase hydrophobicity of the binding pocket. The V287T mutation maintains approximate residue size while introducing a polar side chain to the hydrophobic pocket. The F280Y mutation was previously demonstrated to reduce BBR affinity and was included both as a control for the MD simulation of the PTP1B-BBR complex and to evaluate the effect of this mutation on AD.

This initial list of F196A, L192A, L192F, L192N, L195A, L195F, L195N, F280Y, S286A, E276F, K279M, and V287T were generated using Modeller and we carried out MD simulations using the protocol discussed above with PTP1B initialized with an open WPD loop and disordered *α*7 helix. The MD simulations of the mutant-ligand complexes were examined to determine differences in ligand binding conformation as well as changes in the disruption of inter-helical interactions and the h-bond network. Guided by the MD simulations (more details in section S1.4), a subset of mutations were chosen—F196A, L192F, L195F, E276F, V287T, and F280Y—to examine in experimental studies. These mutations demonstrated distinct differences which we predicted would alter allosteric inhibition in distinct and significant degrees from one another.

For each of the mutations the relative binding free energy was estimated using alchemical transformations via MD simulation. The hybrid topology for each mutation was generated using the PMX web-server and the hybrid residues were parameterized with the PMX hybrid force field.^45^ GROMACS free energy simulations were run utilizing Hamiltonian replica exchange with all parameters available in the “data/mdp” repository folder. The choice of individual alchemical intermediate states were optimized with a single mutation (F196A) in order to maximize overlap between adjacent states, minimize error, and converge estimates from various estimators (Figure S14). These λ states were then used for all other mutations. Analysis of the simulations was completed using alchemlyb 0.6.0^46^ with the TI, MBAR, and BAR estimators. For the final binding free energy estimate, the difference between the change in free energy for PTP1B in solvent and PTP1B-AD complex in solvent was computed and the errors of each estimate were added together (Table S6). The free energy estimates reported are from the MBAR estimator and the error reported for the relative free energy estimates is the analytical error estimated from the sub-sampled 50 ns trajectories at 18 λ states.^47^

### Enzyme Kinetics

We characterized PTP1B activity on p-nitrophenyl phosphate (pNPP) by monitoring the formation of p-nitrophenol (absorbance at 405 nm) in 10-second intervals for 5 min (SpectraMax iD3 plate reader). The composition of each reaction was as follows: PTP (0.05 *μ*M), pNPP (10 mM), BBR (0, 9.6, or 14.4 *μ*M), AD (0, 77, or 115.5 *μ*M), and (50 mM HEPES, 10% DMSO, 50 *μ*g/ml BSA, pH 7.3). The enzyme and inhibitor were incubated together for 5 min before adding the substrate. For each inhibitor, we used previously reported *IC*_50_s^11,33^ to identify the concentrations of each inhibitor that inhibit the wild-type enzyme to a similar extent for both inhibitors (Figure S1). We report kinetic measurements in Table S4.

For each mutant, we evaluated the fractional change in inhibition (F) by using Eq. 1, where *V_o-mut_* and *V_o-wt_* are the uninhibited initial rates of the mutant and wild-type enzyme,

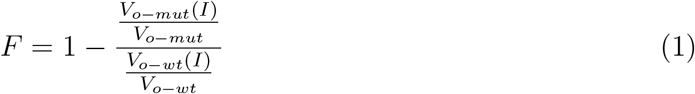

respectively, and *V_o-mut_*(*I*) and *V_o-wt_*(*I*) are the inhibited initial rates. We report values of *V_o-wt_*, *V_o-wt_*(*I*), *V_o-mut_, V_o-mut_*(*I*), and F in Table S4, and we plot values of F in Figure 4B and Figure 5.

**Figure 4:**
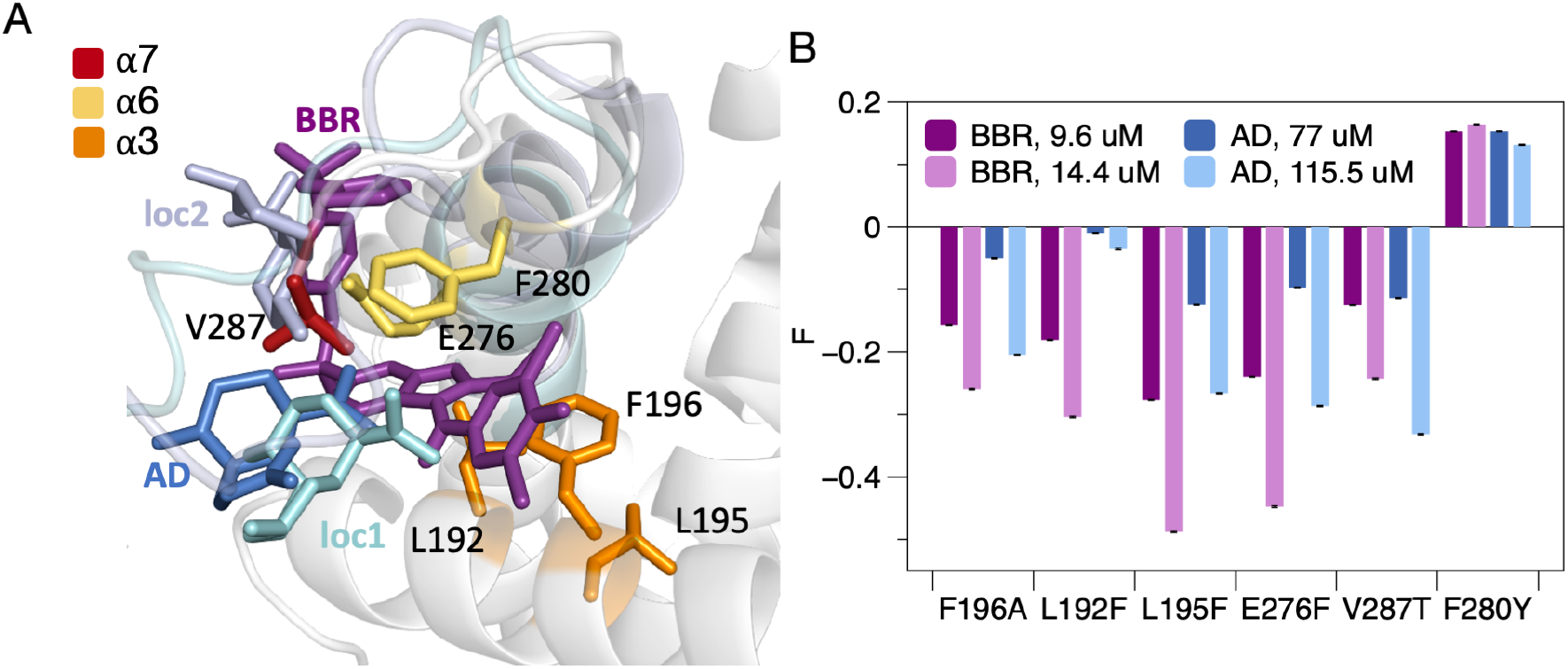
Mutations in the helical triad tend to disrupt inhibition by AD and BBR. (A) An X-ray crystal structure of PTP1B bound to AD (PTPB entry 6W30) with several other binding sites overlaid: (i) the crystallographic binding site for BBR and (ii) two sites sampled by AD in MD simulations (loc 1 and loc2) carried out with a disordered *α*7 helix. To position the alternative sites, we aligned the PTP1B-AD complex (PDB entry 6W30) with the PTP1B-BBR complex (pdb entry 1T4J) and centroid structures from MD simulations (PyMol function “align”). Labels denote residues selected for site-directed mutagenesis with colors by helix. (B) The fractional change in inhibition (F) caused by mutations at the sites highlighted in A. Most mutations decreased the inhibitory effects of AD and BBR. Error bars denote propagated standard error for n=4 independent measurements.

**Figure 5:**
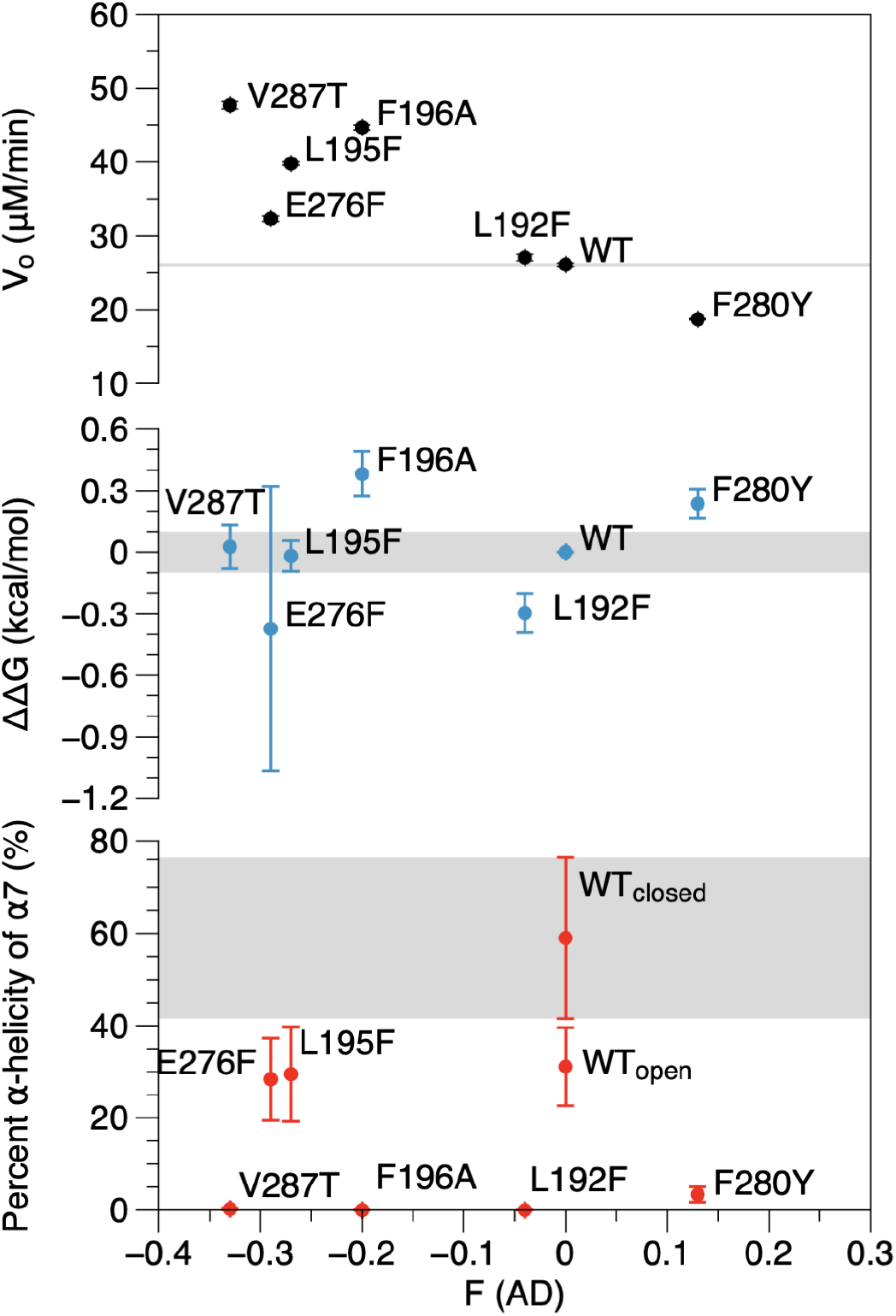
Mutational effects arise from delocalized structural changes in PTP1B. The influence of mutations on AD-mediated inhibition is weakly correlated with their effect on enzyme activity (measured *V_o_*, the initial rate of pNPP hydrolysis in the absence of inhibitor) but not binding affinity (ΔΔ*G*, the difference in free energy of binding between mutants, as calculated from relative free energy simulations using MBAR for analysis) or the mean percent *α* helicity of the *α*7 helix. Shaded regions correspond to the wild-type activity of PTP1B (top), ± 0.1 kcal/mol (middle), and the percent *α* helicity of the wild-type enzyme with WPD_open_ (bottom). Error bars denote standard error for (top) *n* > 4 independent measurements and (middle and bottom) 50 ns simulations at 18 alchemical states. See Materials and Methods for a detailed description of our calculation of standard error for alchemical free energy calculations.

## Results and Discussion

### AD and BBR Bind to Distinct Sites on PTP1B

X-ray crystal structures of PTP1B bound to AD and BBR, a well-characterized benzobromarone derivative,^23^ indicate that they bind to non-overlapping regions of the allosteric site (Figure 2A). We used binding isotherms to assess their ability to bind simultaneously. The catalytic domain of PTP1B has six tryptophan residues that can undergo fluorescence quenching when it binds to ligands. In a preliminary analysis, BBR and TCS401, a well-studied competitive inhibitor, produced strong quenching, while AD had a comparatively small effect—an early indication that it has an unusual binding mode (Table S3). To examine competition between BBR and AD, we measured changes in tryptophan fluorescence caused by BBR in the presence and absence of AD or TCS401 (Figure 2B). TCS401 served as a positive control for orthogonal binding; this ligand causes the WPD loop to close and, thus, binds in a mutually exclusive manner to BBR, which causes it to open. To ensure similar levels of binding by AD and TCS401, we used concentrations that produced similar levels of inhibition (~50%; Figure S1). As expected, AD had a nearly imperceptible effect on the binding isotherm for BBR; the maximum change in fluorescence (Δ*F_max_*), an indicator of the net achievable conformational change, decreased slightly, and *K_d_* remained unchanged. By contrast, TCS401 reduced Δ*F_max_* by over 50%, a change consistent with a reduction in binding sites, and increased *K_d_* by three-fold. This higher *K_d_* suggests that BBR displaces some TCS401, as PTP1B-TCS401 dissociation should have a free energetic penalty; the failure of AD to cause such a change, in turn, suggests that BBR does not displace AD. In general, the insensitivity of the BBR binding isotherm to AD—relative to its extreme sensitivity to TCS401—suggests that AD does not disrupt the binding of BBR.

AD is an unusual inhibitor because it is a hydrocarbon that lacks polar anchoring groups, such as h-bond donors or acceptors (Figure 2A). We speculated that AD might destabilize PTP1B by acting as a nonpolar denaturant—that is, by reducing the free energetic cost of exposing buried residues in water.^48^ Notably, Ertiprotafib, an inhibitor that entered clinical trials, reduces the melting temperature of PTP1B and induces aggregation, an effect consistent with protein denaturation.^49^ We used differential scanning fluorimetry (DSF) to compare the impact of four inhibitors on the stability of PTP1B: AD, BBR, Ertiprotafib (a positive control), and TCS401 (Figure 2C). To our surprise, BBR and Ertiprotafib reduced the melting temperature in a concentration-dependent manner (Δ*T_m_* > 5° C), while AD and TCS401 had no effect (Δ*T_m_* < 1° C, a threshold consistent with prior work^50^). This data indicates that AD does not inhibit PTP1B through nonspecific destabilization.

### AD Stably Samples Two Neighboring Sites on PTP1B

We used MD simulations to study the mechanism by which AD forms a stable complex with PTP1B. To begin, we initialized PTP1B with a disordered *α*7 helix and positioned AD at the crystallographic binding site (loc1). Initially, we used four versions of the disordered *α*7 helix, but only the two conformations with less binding site flexibility (i.e., the two conformations for which residues in the partially disordered *α*7 exhibited lower RMSFs) allowed AD to remain bound during the entire 300 ns trajectory (Figure S4A), with other structures resulting in AD leaving the binding site. This finding suggests that AD binding requires a partially—but not fully—disordered *α*7 helix. We used the two *α*7 conformations that retained AD in the allosteric binding site to model the disordered helix when bound to AD for the remainder of our study.

Over the course of our MD simulations with a disordered *α*7 helix, AD sampled two neighboring sites—loc1 and loc2—with similar occupancies regardless of initial WPD loop conformation (Figure 3). Transitions between the two neighboring sites were infrequent, but observable within the timescale of our simulations, with round trip time from loc1 to loc2 back to loc1 of 55±51 ns, while starting from loc2 the round trip time was was 54±23 ns (more details regarding calculation in section S1.6). The large variance in transition time prevent a direct comparison to binding affinity; however, the similar occupancies suggest the two sites do not have significantly different binding affinities. When we repeated our simulations with an ordered *α*7 helix, or with this helix completely removed, AD sampled two alternative sites located outside of the helical triad: loc3 and loc4, the former had a higher occupancy (Figure 3B; Figure S5). Interestingly, loc3 overlaps with the L16 site, a proposed extension of the allosteric region; however, our simulations indicate binding to either loc3 or loc4 does not produce the same structural changes in PTP1B caused by binding to loc1 and loc2.^25^ Notably, binding to these alternate locations does not disrupt the h-bond network necessary for allosteric inhibition; however, these alternate binding locations are able to increase *α*7 helix disordering when the helix is present. Overall, the importance of a flexible *α*7 helix in facilitating the binding of AD to its crystallographic site (loc1) and a close neighboring site (loc2) is consistent with previous kinetic measurements, which show that the removal of *α*7 reduces the potency of AD.^33^ The contribution of the *α*7 helix may also explain the selectivity of AD for PTP1B over TCPTP. Structural studies suggest that this helix has a similar allosteric function in TCPTP and PTP1B, so differences in inhibition by allosteric inhibitors probably result from differences in ligand binding.^34,51^ Additional biophysical analyses are necessary to determine how the composition and structure of the *α*7 contribute to the selectivity of AD.

To identify unique characteristics of the PTP1B-AD interaction, we also used MD simulations to study the binding of BBR. We initialized PTP1B with both an ordered and disordered *α*7 helix, as prior kinetic experiments indicate that this helix enhances inhibition by BBR.^22^ Curiously, in a subset of trajectories carried out with the ordered *α*7 helix, BBR bound with an elongated conformation that differed from its crystallographic pose (Figure S7); we focused our analysis on *α*7 conformations that allowed BBR to bind with a conformation consistent with the crystal structure. Like AD, BBR exhibited prominent interactions with the helices of the helical triad, including *π*-stacking with F280. BBR engaged in more interactions with the *α*7 helix; AD, the *α*3 helix (Figure 3D; Figure S8). Unlike AD, BBR did not oscillate between different binding locations, a behavior demonstrated by higher RMSF of the ligand COM around the ligand centroid for AD compared to BBR (Figure 3E).

### AD Moves to Accommodate BBR in a Ternary Complex

Guided by experimental evidence that AD and BBR bind to different sites, we used molecular simulations to examine the simultaneous binding of AD and BBR. We initialized PTP1B with a disordered *α*7 helix and positioned AD and BBR at their non-overlapping crystallographic binding sites. To our surprise, BBR displaced AD, which moved quickly (~500 ps) to a patch formed by residues 290–295 on the outside of the *α*7 helix, about 8.5 Å from its initial binding site. AD stayed at this position for the remainder of the 1 *μ*s simulation (Figure 3C). In the ternary complex, the flexibility of the region of *α*7 in contact with AD decreased, while the rest of the helix became more flexible (relative to the PTP1B-AD or PTP1B-BBR complexes; Figure S4B). The flexibility of AD also decreased, a change in mobility consistent with stabilization by BBR (Figure 3E). The ability of AD and BBR to bind simultaneously to PTP1B is consistent with our binding data (Figure 2B).

### PTP1B Mutations Elicit Delocalized Structural Changes

MD simulations and kinetic analyses allowed us to find mutations that disrupt inhibition by BBR and AD to different extents, and facilitated a detailed analysis of the effect of structural perturbations on the allosteric site. As described in the methods, we began by identifying residues that modify the binding pose of AD or BBR or that disrupt the helical triad in MD simulations; we used site-directed mutagenesis to change the size or chemical functionality of these residues; and we used in vitro kinetic assays to measure the change in inhibition caused by each mutation (Figure 4B). Importantly, for this analysis, we used concentrations of AD and BBR that inhibit the wild-type enzyme to the same extent—a constraint that enables comparisons of the fractional change in inhibition between compounds (Figure S1). All but one mutation (F280Y) reduced inhibition by both inhibitors and, in general, mutational effects were more pronounced for BBR than for AD. The reduced sensitivity of AD is consistent with its ability to adopt multiple bound conformations.

A comprehensive analysis of the influence of mutations on enzyme activity, binding affinity, and *α*7 helix structure suggests that they do not disrupt allosteric inhibition by impeding protein-ligand binding. Several apo mutants exhibited differences in catalytic activity. For AD, these differences were weakly correlated with changes in inhibition (Figure 5); this relationship suggests that the mutations, which sit outside of the active site, might nonetheless affect communication with that site. We used relative free energy simulations to estimate mutant-derived changes in binding free energy. These changes were not correlated with inhibition (Figure 5). Given the current limitations on the accuracy of the relative free energy protocol, which provides a maximum resolution of ~1 kcal/mol,^52^ a direct comparison between binding affinities is challenging; nonetheless, our analysis suggests that the mutations do not cause a statistically significant change in binding affinity. MD simulations allowed us to examine the influence of mutations on the *α* helicity of the *α*7 helix, an important constituent of the allosteric site. Most mutations destabilized this helix, but this change was not predictive of their influence on inhibition (Figure 5).

Analysis of the helical triad suggests that delocalized structural changes caused by the mutations contribute to both differences in the level of inhibition caused by both inhibitors and changes in the catalytic activity of PTP1B. Subtle structural changes to the *α*3 helix affected WPD loop movement by limiting the structural changes necessary to reorient it from an open to closed conformation. Changes to the *α*6 helix could potentially alter the allosteric network, but statistically significant conclusions were difficult to make (Figure S5 and section S1.5). In the end, the molecular basis of specific mutational effects proved difficult to resolve in fine detail, but our biophysical analyses, taken together, indicate that mutations disrupt inhibition through delocalized structural changes—that is, through small structural changes that do not significantly affect ligand binding but, rather, the allosterically influential conformational changes that result from that binding.

### AD and BBR Destabilize the *α*7 Helix and Disrupt WPD Loop Motions

We sought to determine how AD modulates PTP1B activity by using MD simulations to trace allosteric communication between its binding site and the active site. We began by mapping the h-bonds that link ordering of the *α*7 helix and closure of the WPD loop (Figure 6B; Figure S10). These bonds connect (i) the WPD to the P-loop, (ii) the WPD-loop to the L-11 loop and the *α*6 helix, and (iii) the L-11 loop to the *α*3 and *α*7 helices (Figure 6A). This set of regions match those identified in prior work, but two of the connecting h-bonds are new^18,22,53^ (Figure S11). An additional bond connecting the L-11 loop directly to the *α*7 helix (Y152–S295) as well as an additional bond connecting the WPD-loop and *α*6 helix (T263–F182) are unique to this analysis. These additional bonds suggest that allosteric communication within PTP1B may include some redundancy—that is, different h-bonds may permit communication between the same neighboring sites. Simulations with a closed WPD loop and a disordered *α*7 helix revealed that helix disordering alone disrupts the formation of the bonds N193–Y152 and N193–E297; the remainder of the bonds are disrupted via the reorientation of the WPD loop (Figure S12). This suggests that the prevention of *α*7 ordering should prevent the h-bond network from forming and thus prevent the closure of the WPD loop (Figure 6A–B).

**Figure 6:**
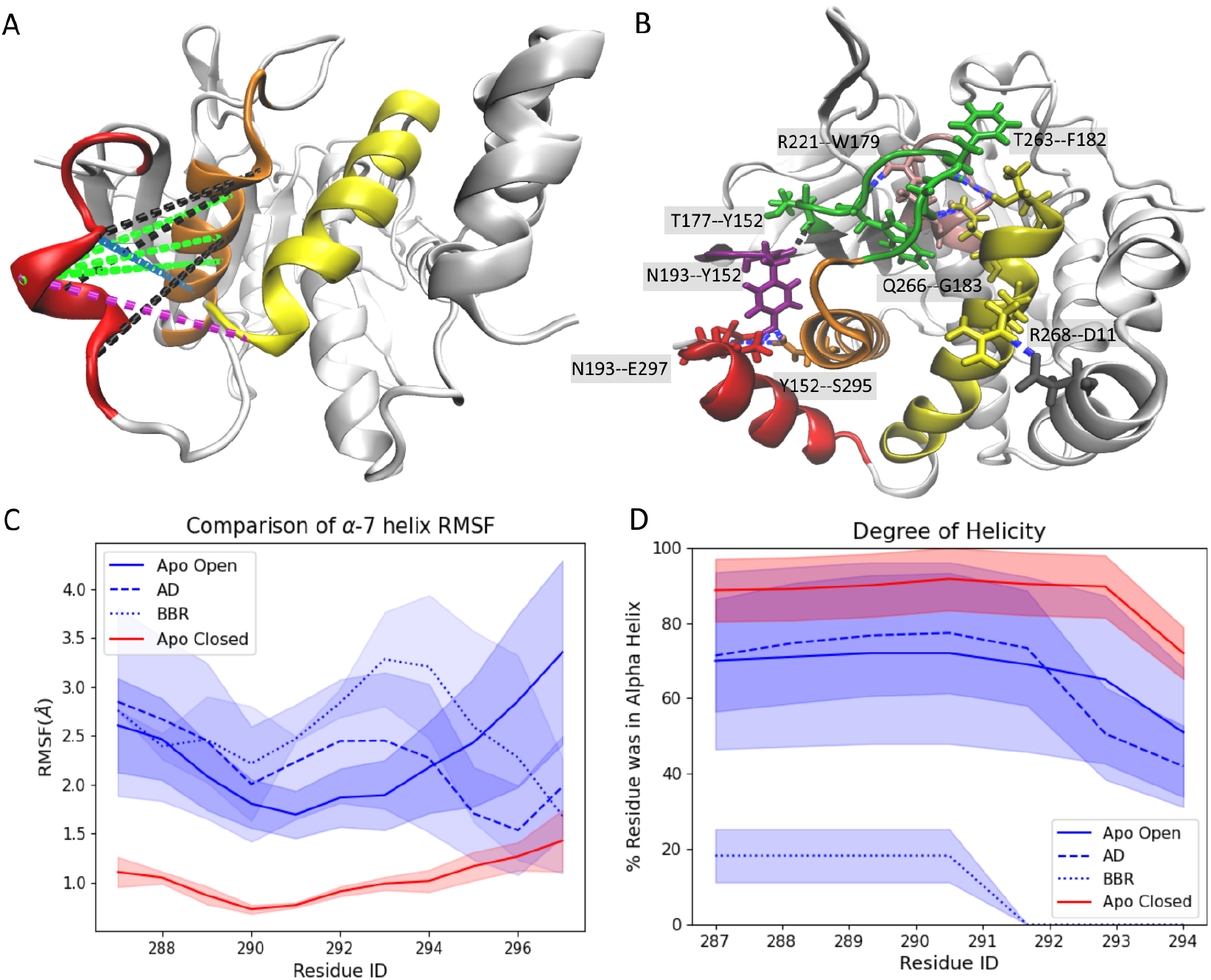
Ligand binding disrupts inter-helical interactions. (A) Upon binding, AD and BBR disrupt non-bonded interactions between the *α*3, *α*6, and *α*7 helices. Highlights: (black) interactions disrupted in the WPD*_open_* conformation that are also disrupted when AD and BBR bind to the protein, (green) interactions disrupted by both ligands but present in both WPD_closed_ and WPD_open_ conformations, (purple) interactions disrupted by BBR alone, and (blue) interactions disrupted by AD alone (Figure S6). Most of the interactions disrupted by AD, BBR, and WPD*_open_* are located between the *α*3 and *α*7 helices. This overlap suggests that that the disruption of these interactions is crucial for allosteric inhibition. (B) Disruption of the interactions depicted in A destabilizes the *α*7 helix and prevents the formation of h-bonds required for closure of the WPD loop. The blue lines denote bonds that form with WPD_closed_ but don’t form with apo WPD_open_ and are broken with ligand binding. The black line denotes the conserved bond T177–Y152 which connects the active and allosteric sites. (C–D) Upon binding, AD (dashed line) and BBR (dotted line) (C) increase the flexibility of the *α*7 helix and (D) decrease its *α* helicity to levels that resemble the *WPD_open_* conformation. Destabilization of the *α*7 helix is faster with BBR than with AD (Figure S9). In C–D, all MD trajectories start with the same ordered *α*7 helix conformation.

We evaluated the influence of AD and BBR on *α*7 stability by initializing MD simulations with an ordered helix. Both inhibitors enhanced the flexibility of the *α*7 helix and accelerated its disordering, relative to the apo structure (Figure 6C–D). For the ligand-bound states, the initial WPD loop conformation did not affect helix disordering; by contrast, when we initialized the Apo state with an open WPD loop, helix disordering increased relative to simulations initialized with a closed WPD loop. Intriguingly, AD enhanced helix disordering while bound to its alternative sites (loc3 and loc4) and may shift back to loc1 and loc2 when the helix becomes sufficiently disordered, a shift that would likely take too long for our 300 ns simulations to capture.

Our simulations suggest that AD, like BBR, prevents reordering of the *α*7 helix and WPD loop closure by disrupting interactions between the *α*3 and *α*7 helices (Figure 6A; Figure S13). For ligand-bound simulations, the disrupted interactions within the helical triad were conserved for both disordered *α*7 helix conformations regardless of the initial WPD-loop conformation. Apo simulations were initialized with structures consistent with experimental observation: an open WPD loop accompanied a disordered *α*7 helix; a closed WPD loop, an ordered *α*7 helix. When compared with the open-and-close motions of the WPD loop, localized changes in protein conformation, specifically changes in the helical triad, equilibrate quickly and are easier to observe within the time scale of these simulations. In the apo enzyme, the WPD_open_ conformation lacks *α*3–α7 interactions that form when the WPD loop is closed; upon binding, AD and BBR disrupt additional interactions between these two helices (Figure S13A). The disrupted interactions are largely isolated to residues 188–196 on the *α*3 helix and 289–291 on the *α*7 helix (Figure S6). BBR also disrupts *α*6–α7 interactions, but AD does not (Figure S13C). Accordingly, this effect appears non-essential for allosteric inhibition. Changes in *α*3–α7 interactions, by contrast, are conserved between the two inhibitors, an indication that they are centrally important to allosteric inhibition.

## Conclusions

Unfunctionalized terpenoids are a surprising source of selective inhibitors because their nonpolar structures seem well suited to engage in nonspecific interactions with nonpolar regions on protein surfaces.^54^ In this study, we examined the inhibition of PTP1B by AD, a surprisingly potent and selective inhibitor for a small, 15 carbon hydrocarbon. MD simulations indicate that AD samples two adjacent sites near the C-terminus of the catalytic domain, both of which require a disordered *α*7 helix. Intriguingly, when the helix is fully ordered, AD binds to two alternative sites and destabilizes the helix. Perhaps most importantly for inhibitor design, the binding mode of AD is distinct from BBR, a well-studied allosteric inhibitor. DSF data indicate that AD does not destabilize the protein, and binding data and MD simulations suggest that the two molecules can bind simultaneously. Efforts to bridge these two molecules—or the distinct binding sites that they adopt in the ternary complex—could yield more potent inhibitors of PTP1B.

The inhibitory mechanisms of AD and BBR are similar, but the two inhibitors do not induce all of the same conformational changes to PTP1B. MD simulations of apo PTP1B show a network of h-bonds that link ordering of the *α*7 helix to closure of the WPD loop. The sections of PTP1B connected by this network match those uncovered in prior work, but our network contains additional h-bonds, an indication that allosteric communication within PTP1B may have some redundancy. Upon binding, both AD and BBR destabilize the *α*7 helix and prevent closure of the WPD loop. The two inhibitors cause distinct structural perturbations, but both disrupt the *α*3–α7 interface, an indication that this inter-helical interaction is centrally important to allosteric inhibition. The concentration of disrupted interactions at this interface is likely responsible for the disordering of the *α*7 helix and disruption of h-bond network that ultimately reduces catalytic activity. Though disruption of the *α*6–α7 interface has been reported as important to allosteric inhibition, the behavior of AD suggests it is not essential. Our findings also indicate that mutations in the helical triad, defined by *α*3, *α*6, and *α*7, can weaken the effects of AD and BBR, likely by affecting communication with the active site, rather than ligand binding. Similar mutations might confer resistance to therapeutics that rely on allosteric inhibition.

Terpenoids are the largest class of natural products, but many—if not, most—are decidedly non-druglike, at least in their underivatized form.^55^ As a result, they tend to be overlooked at the earliest stages of drug discovery.^56^ This analysis provides evidence that unfunctionalized terpenoids can engage in specific interactions with protein surfaces, and it provides an interesting model system for studying these interactions. By elucidating novel binding modes or allosteric mechanisms, such as those exhibited by AD, protein-terpenoid interactions could inform the design of new varieties of enzyme inhibitors.

## Supplemental Information

Supplementary figures detailing analyses of alternative conformations of the *α*7 helix, relative free energy calculations, binding-induced changes in WPD loop conformation, interactions between AD and BBR and the helical triad, allosteric communication within PTP1B, differential scanning fluorometry, and AD- and BBR-mediated inhibition of PTP1B; tables describing the binding conformations of AD in MD simulations, primers used for site-directed mutagenesis, initialization criteria used for MD simulations, binding measurements, and kinetic measurements.

## Accession Codes

UniProt Accession IDs: PTP1B (P18031) and TCPTP (P17706).

## Author Contributions

A.J.F, J.M.F. and M.R.S conceptualized the project. A.J.F. and M.R.S. designed the computational methodology, and E.T.L, L.K., A.S., and J.M.F. designed the experimental methodology. A.J.F. performed and analyzed all molecular simulations experiments. A.J.F. wrote the original draft. J.M.F. and M.R.S. edited and reviewed the manuscript. E.T.L carried out site-saturation mutagenesis, protein purification, and inhibition measurements. L.K. performed differential scanning fluorimetry experiments. A.S. completed binding studies. J.M.F. and M.R.S. supervised the project and obtained the resources.

## Acknowledgements

This work was supported by funds provided by the University of Colorado Boulder (A.J.F and M.S.), the National Institute of General Medical Sciences of the National Institutes of Health (E.T.L., R35GM143089), the National Science Foundation (A.S. and J.M.F., CBET 1750244), and Think Bioscience (L.K.). This work utilized computational resources from the University of Colorado Boulder Research Computing Group, which is supported by the National Science Foundation (awards ACI-1532235 and ACI-1532236), the University of Colorado Boulder, and Colorado State University. This work also used the Extreme Science and Engineering Discovery Environment (XSEDE), which is supported by National Science Foundation grant number ACI-1548562. Specifically, it used the Bridges-2 system, which is supported by NSF ACI-1928147 at the Pittsburgh Supercomputing Center (PSC).

## Abbreviations

PTP1B: protein tyrosine phosphatase 1B
TCPTP: T-cell protein tyrosine phosphatase
MD: molecular dynamics
AD: amorphadiene
BBR: 3-(3,5-Dibromo-4-hydroxy-benzoyl)-2-ethyl-benzofuran-6-sulfonicacid-(4-(thiazol-2-ylsulfamyl)-phenyl)-amide
TCS401: 2-[(Carboxycarbonyl)amino]-4,5,6,7-tetrahydro-thieno[2,3-c]pyridine-3-carboxylic acid hydrochlo-ride
DSSP: Defined Secondary Structure Prediction

