## Supplementary Information for "Allosteric Inhibition of PTP1B by a Nonpolar Terpenoid"

### Supporting Information: Allosteric Inhibition of PTP1B by a Nonpolar Terpenoid

#### S1 Methods Continuation

##### S1.1 Correlated Frames Removal

Correlated frames were removed using ruptures. Ruptures is a change point detection software that allows for segmentation of non-stationary signals like trajectory RMSD values based on change points. We determined change points from the trajectory backbone RMSD using the binary segmentation method with a threshold on the residual norm. Uncorrelated frames were defined as those in which change points were detected. The trajectory was then limited to these uncorrelated frames for all further analysis. The equilibrated and uncorrelated trajectories included 409$\pm$59 configurations which is one configuration every $\sim$0.7 ns. The removal of these uncorrelated trajectory frames reduce the likelihood of over-estimating statistical significance of time dependent measurements.

##### S1.2 Equilibration

We defined the equilibration point as the point at which the backbone RMSD fluctuations of the trajectory plateau and the mean of 100 ps simulation blocks does not vary by more than 0.5 Å consistently for a minimum of 1 ns. The equilibration time was then rounded up to the nearest 5ns interval. Following equilibration of the simulation trajectory, we tested to ensure the simulation had converged at least in regards to the protein backbone. Convergence was defined as a difference in RMSD between adjacent frames of less than 0.2 Å for the full equilibrated trajectory. Frames prior to equilibration were removed from analysis unless when explicitly stated. The mean time to reach equilibration was 47$\pm$6 ns for all trajectories with a mode equilibration time of 35 ns.

##### S1.3 Ligand RMSD

We clustered the simulations with ligands bound on the ligand heavy atoms. We concatenated all trajectories with the same ligand bound in the crystallographic binding pose prior to clustering. These trajectories had initial conformations with the WPD loop both open and closed and a disordered $\alpha7$ helix which resulted in AD bound to either loc1 or loc2 and BBR bound in the crystallographic pose. The centroid of the most populated cluster was taken as the ligand centroid. We computed the RMSD for the ligand center of mass (COM) relative to these ligand centroid structures. We determined the COM coordinates using the `MDTraj compute_center_of_mass` and computed the RMSD using the following equation:

$$RMSD = \sqrt{\frac{1}{T}\sum_{t=0}^{T}(x(t) - x_{ref})^2 + (y(t) - y_{ref})^2 + (z(t) - z_{ref})^2} \ .$$

Bootstrapping was then performed on the full ligand COM RMSD for the concatenated trajectory. Each iteration randomly selected 200 samples from the RMSD values. A total of 20 iterations were performed for each concatenated trajectory. The same protocol was performed for the ternary complex trajectory separately for both AD and BBR; however, since there was only one trajectory there was no need to concatenate trajectories.

#### S1.4 Mutant Selection

The initial list of mutations which were simulated was reduced to the mutations which showed the highest likelihood for conformational changes which would alter inhibition of PTP1B by AD, BBR, or both. Overall significant differences in the h-bond network were not observed except in the case that the ligand dissociated and the WPD loop reoriented to a closed conformation. There were, however, significant differences in ligand placement within the binding pocket which resulted in significant differences in the ligand disruption of interactions in the helical triad. F196A-AD complex showed a decrease in the disruption of interactions at the $\alpha3$–$\alpha7$ helix interface and an increase in the disruption of interactions at the $\alpha6$–$\alpha7$ helix interface. Meanwhile the F196A-BBR complex showed a reduction in the interaction disruption with both helices. If our prediction that the disruption of the $\alpha3$–$\alpha7$ helix interface is most significant than both ligands would show decreased inhibition in experimental observation. The L192F-AD complex shows no significant change in the $\alpha3$–$\alpha7$ helix interface while the L192F-BBR complex shows increased disruption indicating that inhibition by BBR may increase while inhibition by AD would be relatively unaffected. The opposite was observed with the L195F-AD and L195F-BBR complexes. E276F showed similar changes as L195F mutation but with the addition of a lack of disruption of interactions in the $\alpha6$–$\alpha7$ helix interface for the E276F-BBR complex. V287T-AD complex showed an increase in the disruption of interactions at both the $\alpha3$–$\alpha7$ helix and $\alpha6$–$\alpha7$ helix while the V287T-BBR complex showed a reduction in the disruption of interactions at the $\alpha3$–$\alpha7$ helix and increase at the $\alpha6$–$\alpha7$ helix. F280Y-AD and F280Y-BBR complexes were both unstable in MD simulations suggesting a reduction in binding affinity which would produce decreased inhibition by both AD and BBR. Additionally mutants differed in the interactions which were disrupted at the helical interfaces. The mutations which were not selected either showed the same differences as those selected but with a smaller magnitude of change or showed no significant differences from the WT-ligand complexes.

#### S1.5 Delocalized mutation effects

We probed the structural basis of mutational effects further by examining catalytically influential regions of PTP1B. Mutations L192F, L195F, E276F, and V287T caused residues 186–191, which connect the WPD loop to the $\alpha 3$ helix, to shift their orientations toward those adopted in the $WPD_{closed}$ state (Figure S5A,C). These shifts could affect how interactions with the $\alpha 3$ helix influence WPD loop motions. Mutations L192F, L195F, and E276F, in turn, caused the $\alpha 6$ helix to rearrange, which might also disrupt allosteric communication with the active site, though this rearrangement was not consistent across mutations (Figure S5B,D). Interestingly, all mutations stabilized the catalytic C215 and reoriented the L1 loop (residues 27–35; Figure S5F–G), a common influence uncorrelated with changes in inhibition.

#### S1.6 Determination of AD Occupancy Percentages

In order to determine the percent time that AD occupied the different binding locations we first eliminated the non-physical trajectories. These included any trajectories with a truncated or ordered $\alpha 7$ helix or which maintained a WPD loop in the closed conformation (if the WPD loop was initialized closed but reoriented to an open conformation than the trajectory was kept). Then we removed any trajectories which did not maintain AD bound to PTP1B for a majority of the trajectory (>75%). This left us with three AD bound trajectories, one with the "disordered 3" $\alpha 7$ helix conformation and two with "disordered 4". Disordered conformations 1-3 were generated from the restrained heating method and conformation 4 was generated from the MD simulation of of the PTP1B-BBR complex as described in the materials and methods sections. We make the assumption that the two disordered conformations are equally likely and thus we weigh the trajectories to equalize their contribution to the occupancy percentages. We also computed the occupancy of AD in alternative binding locations. This calculation was similar but we limited to only trajectories with either an ordered or absent WPD loop and no weighting was required since there were no differences in the $\alpha 7$ helix among each group.

For the three trajectories with AD bound to either loc1 or loc2, a transition time was calculated. Only two of the three trajectories featured transitions between the two binding locations. Since these two trajectories only had AD bound in either loc1 or loc2, two different transitions were defined: from loc1 to loc2 to loc1 and reversed. Combining the two trajectories there are four transitions from loc1 to loc2 to loc1 and three from loc2 to loc1 to loc2. The time in nanoseconds was computed for each of the transitions and the mean and standard deviation was calculated.

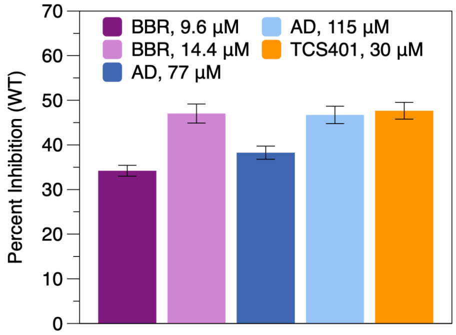


Figure S1: Inhibition of wild-type PTP1B. For our binding study, we used concentrations of AD and TCS401 (115$\mu$M and 30$\mu$M, respectively) that inhibit PTP1B by $\sim$50% to ensure that the two ligands interact with PTP1B to a similar extent. For our mutational analysis, we used two concentrations of BBR and AD that produce similar levels of inhibition for both ligands.

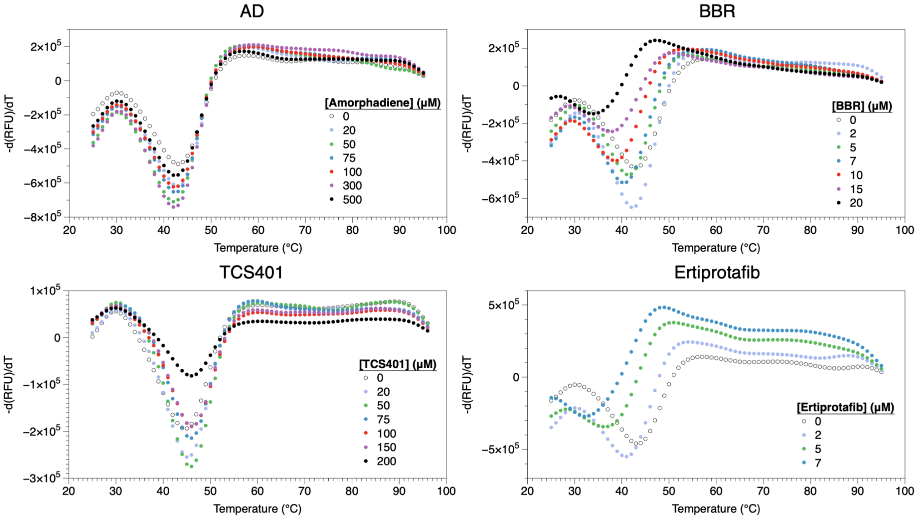


Figure S2: Differential scanning fluorimetry (DSF) of four PTP1B inhibitors. We examined the influence of different inhibitors on the melting temperature ($T_m$) of PTP1B by measuring changes in the fluorescence of SYPRO orange as a function of temperature at different inhibitor concentrations. These plots show the negative of the first derivative of fluorescence vs. temperature. We used the minimum of each curve to calculate $T_m$ at each inhibitor concentration (Fig. 2C)[2]. Note: The shifted baseline and reduced melting temperature caused by BBR and Ertiprotafib suggest that these molecules destabilize PTP1B.

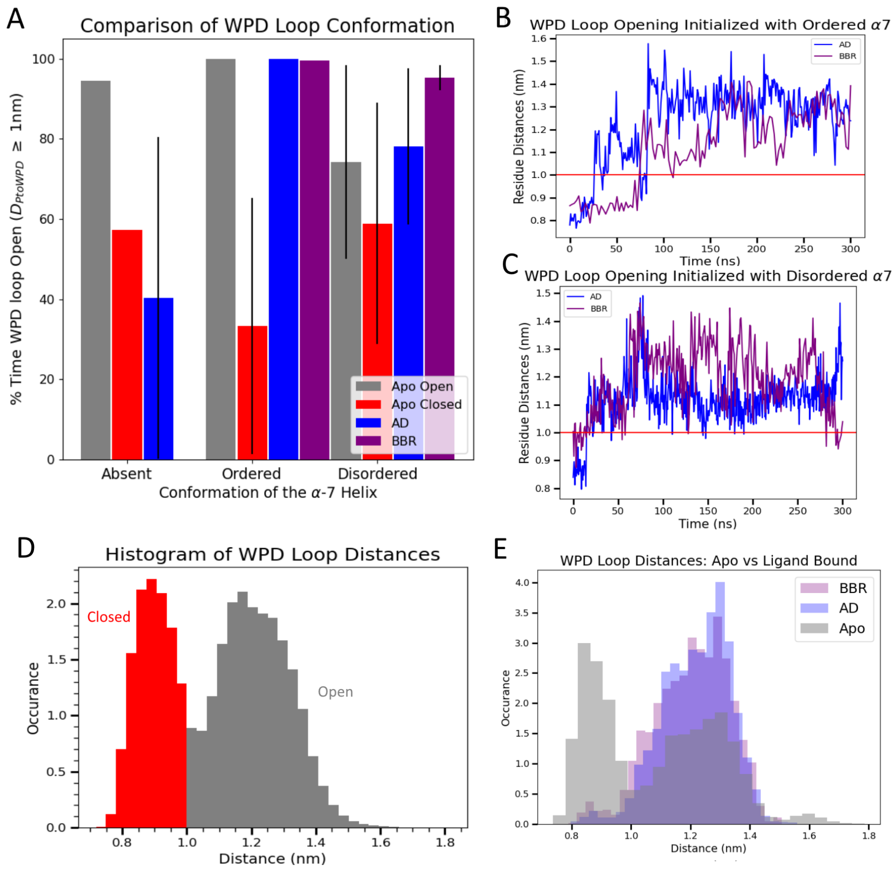


Figure S3: AD and BBR binding to the allosteric site reorient the WPD loop to an open conformation. (A) When we initialized PTP1B with the WPD loop in a closed conformation and AD or BBR positioned at the crystallographic binding site, the WPD loop shifted to an open conformation. This shift occurred regardless of the starting conformation of the $\alpha 7$ helix (i.e., ordered or disordered), but not when this helix was absent. Error bars denote standard error between simulations run with the same initial configurations. The WPD loop reorients into an open conformation more quickly for (B) AD than for BBR and (C) even more so when the protein begins with a disordered rather than ordered $\alpha 7$ helix. (D) A histogram of WPD loop distances over all apo simulations is distinctly bimodal. We used the minimum at $\sim 1$ nm for the cutoff to differentiate between open and closed conformations of the WPD loop. (E) All simulations show a bimodal distribution, but the binding of AD or BBR shifts the distribution towards the open conformation compared to apo simulations.

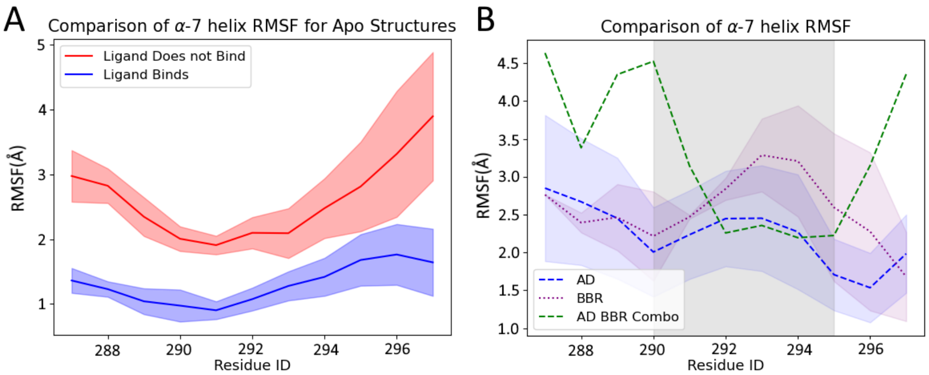


Figure S4: The binding of AD and BBR requires partial, but not complete ordering of the $\alpha7$ helix. (A) The root mean square fluctuations (RMSFs) for four versions of the disordered $\alpha7$ helix used in initial simulations of PTP1B-AD and PTP1B-BBR complexes. The two conformations that retained AD and BBR (blue) were less flexible than those that did not (red). The blue line is the mean RMSF of the 3 apo trajectories (two with the WPD loop initialized open and one closed) corresponding to the disordered helix conformations which produced a stably binding ligand. The red line is the mean RMSF of the 4 apo trajectories (two with the WPD loop initialized open and two closed) corresponding to the helix conformation which did not produce a stably binding ligand. The shaded region for both lines denotes the uncertainty interval which is the standard error of the RMSF of each residue from the corresponding trajectories in which the ligand either does or does not remain stably bound. (B) When bound to an allosteric ligand, the flexibility of the $\alpha7$ helix increases (from 1-3Å to 1.5-3.5Å). This effect is statistically indistinguishable for AD (blue) vs BBR (purple); however, when both ligands are bound simultaneously there is a significant increase in the flexibility of the N- and C- terminal ends of the $\alpha7$ helix, though not where AD binds in the ternary complex (shaded gray). The binding of AD to the outside of the $\alpha7$ helix appears to stabilize the helix motions. The blue and purple shaded regions again represent the standard error of the RMSF between the four trajectories with AD and three trajectories with BBR both bound in their crystallographic pose (or loc2 for AD). No error region is provided for the ternary complex as only one $1\mu s$ trajectory was completed.

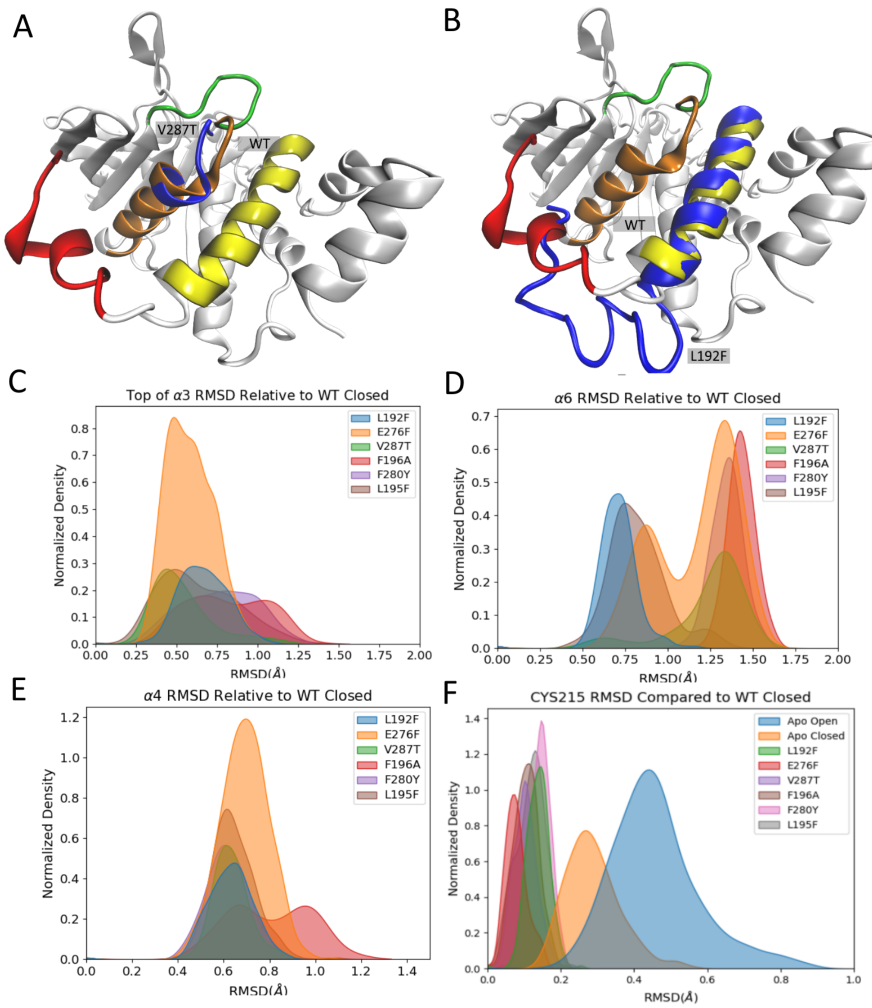


Figure S5: Mutations in the allosteric site caused non-localized conformational changes in PTP1B. (A) The loop between the $\alpha3$ helix and the WPD loop reoriented as the WPD loop transitioned from open to closed states. (C) Several mutants exhibited a conformation of this loop closer to the closed conformation despite retaining an open conformation. (B) Additionally, the $\alpha6$ helix, particularly residues 276–280, reoriented between the open and closed conformations. (D) Mutants L192F, L195F, and E276F adopted conformations closer to the Apo closed state and showed decreased inhibition by both AD and BBR. Opening of the $\alpha6$ helix may disrupt the ligand binding to the appropriate location within the allosteric site and/or limit the ability of the ligand to modulate WPD loop motion once bound. (E) For F196A, the $\alpha4$ helix adopted a distinct conformation with a high RMSD between both the apo open and closed conformations. (F)The catalytic CYS215 residue did not adopt a distinct conformation for any of the mutants evaluated but it had a more stable conformation for all mutants with a more narrow RMSD distribution than either $WPD_{open}$ or $WPD_{closed}$

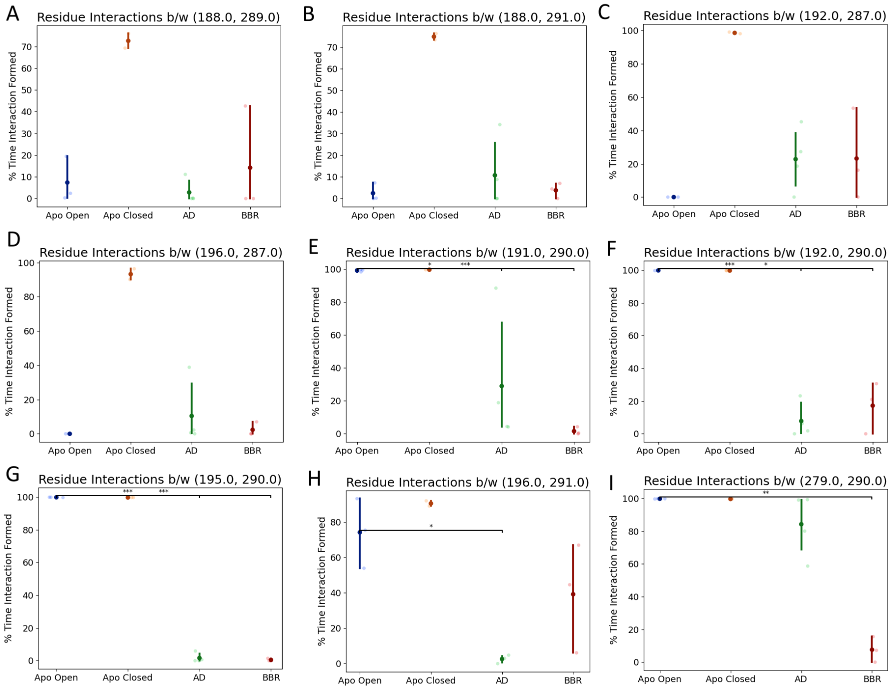


Figure S6: When the WPD loop opens, interactions between residues in the $\alpha3$ and $\alpha7$ helices are consistently disrupted (A–D). The $\alpha3$-$\alpha7$ interactions form for a significantly smaller percentage of the trajectory when the WPD is open than when it is closed. The binding of AD and BBR disrupt additional interactions between these helices (E–G). (H) AD disrupts an interaction between the $\alpha3$ and $\alpha7$ more frequently and consistently than BBR. (I) BBR disrupts an interaction between the $\alpha6$ and $\alpha7$ helices while binding of AD does not.

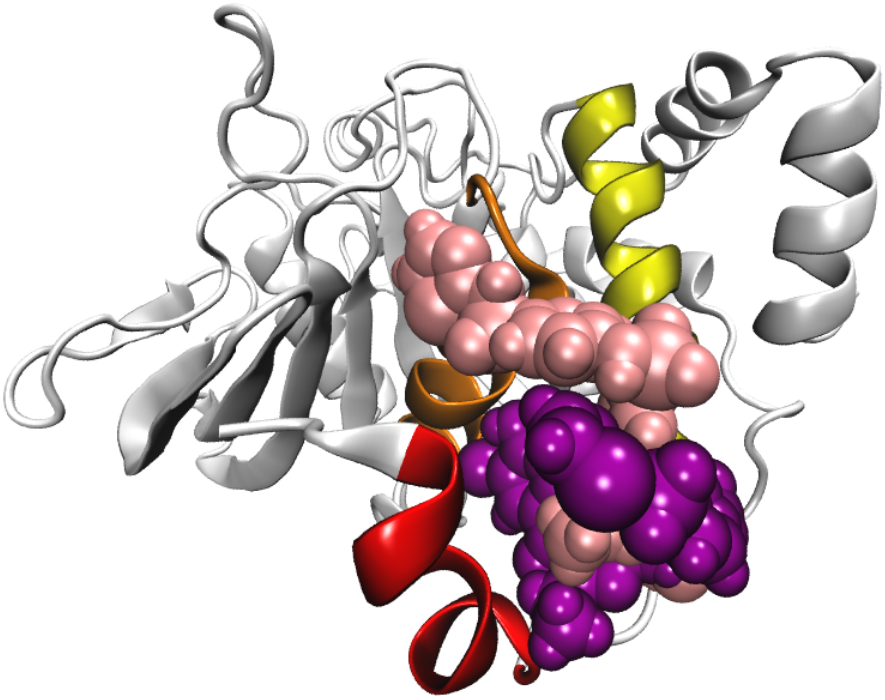

Figure S7: In a subset of the trajectories run with the PTP1B-BBR complex, BBR adopts an alternative conformation (pink) rather than the crystallographic pose (purple). This elongated BBR conformation results in a closed WPD loop, a conformation that is not consistent with biophysical analyses and likely not physically realistic. This alternative binding pose, which occurs with only a subset of disordered and ordered $\alpha 7$ helix conformations, suggests that BBR, like AD, is sensitive to the structure of the $\alpha 7$ helix. Given its larger size, BBR appears less likely to leave the helical triad than AD when the $\alpha 7$ helix conformation is sub-optimal; rather, it adopts an elongated conformation not conducive to inhibition of PTP1B.

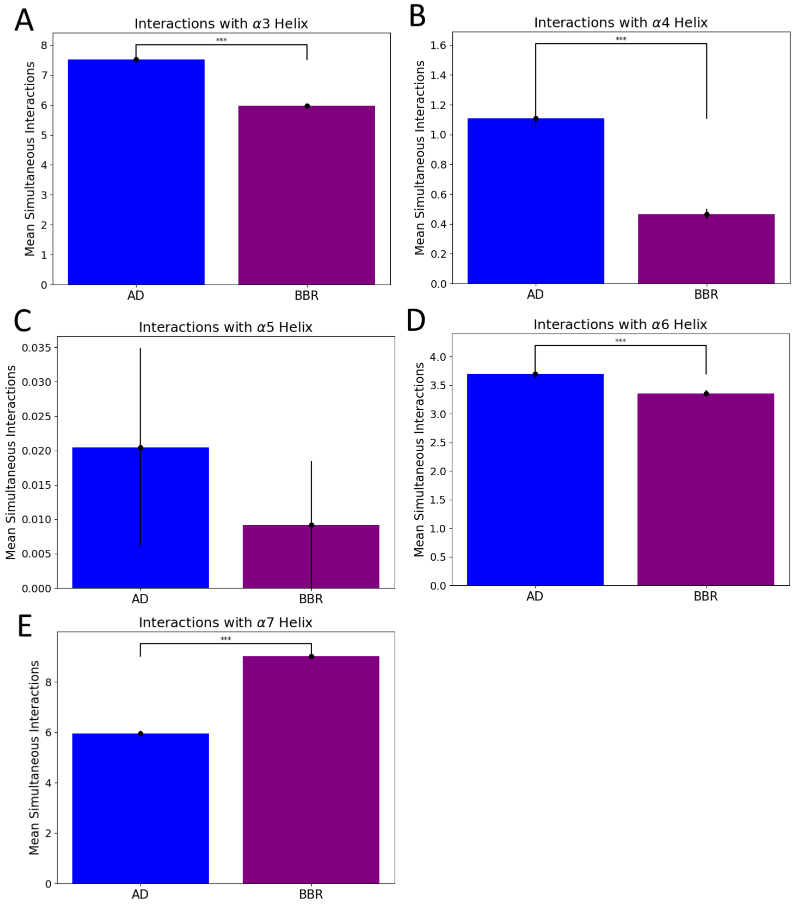


Figure S8: Interactions between AD or BBR and the helical triad are largely similar, but feature some statistically significant distinctions. The number of simultaneous interactions formed between AD and BBR with the $\alpha$4-$\alpha$6 helices were statistically indistinguishable, but the ligands were differentiated by their interactions with the $\alpha$3 and $\alpha$7 helices. Here, we define simultaneous interactions as the total number of residues within the given helix with which the ligand has an interaction ($\leq$ 5 Å). (A) AD formed significantly more (38%) interactions with the $\alpha$3 helix than BBR. (B–C) Neither ligand formed a significant ($\geq$ 1) number of interactions with the $\alpha$4 or $\alpha$5 helices. (D) AD had statistically significantly more interactions with the $\alpha$6 helix, but the magnitude of this difference is small ($<$ 0.5 interaction difference in the mean), suggesting that it may be an artifact of the large number of sampled values. (E) BBR, by contrast, engaged in 18% more simultaneous interactions with the $\alpha$7 than AD—an appreciable difference.

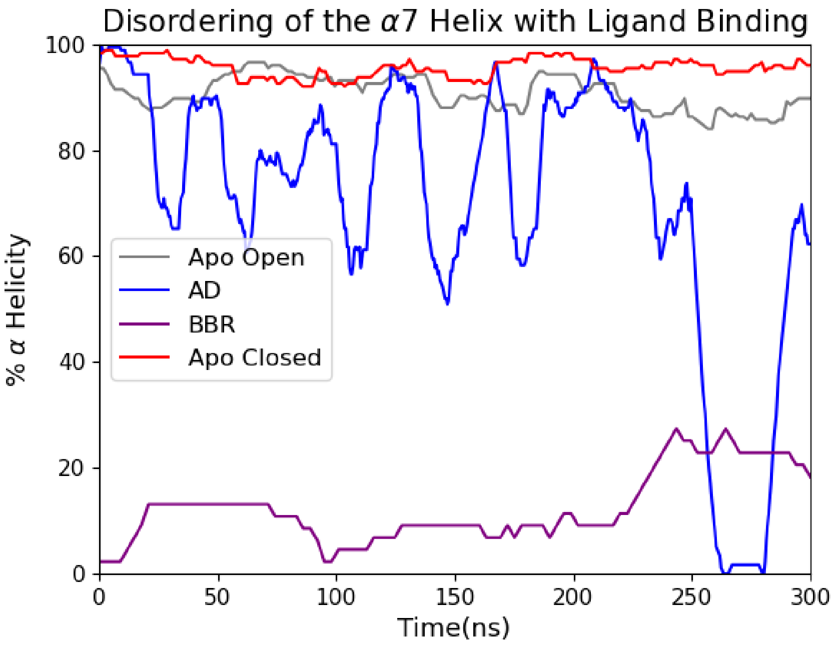


Figure S9: Progressive disordering of the $\alpha7$ helix occurs when either AD or BBR are bound to PTP1B. The $\alpha7$ helix quickly disorders upon binding of BBR to the allosteric site. This disordering occurs during the equilibration period for the PTP1B-BBR complex and is thus not included in the graph. The disordering of the $\alpha7$ helix due to AD binding to the allosteric site is delayed compared to BBR binding. We hypothesize that this delay is due to AD binding to an alternative binding location (loc3 or loc4) outside of the helical triad. It is possible that, given sufficient time, AD would reorient to the crystallographic binding pose (loc1 or loc2) although this was not captured in the time scale of our simulations.

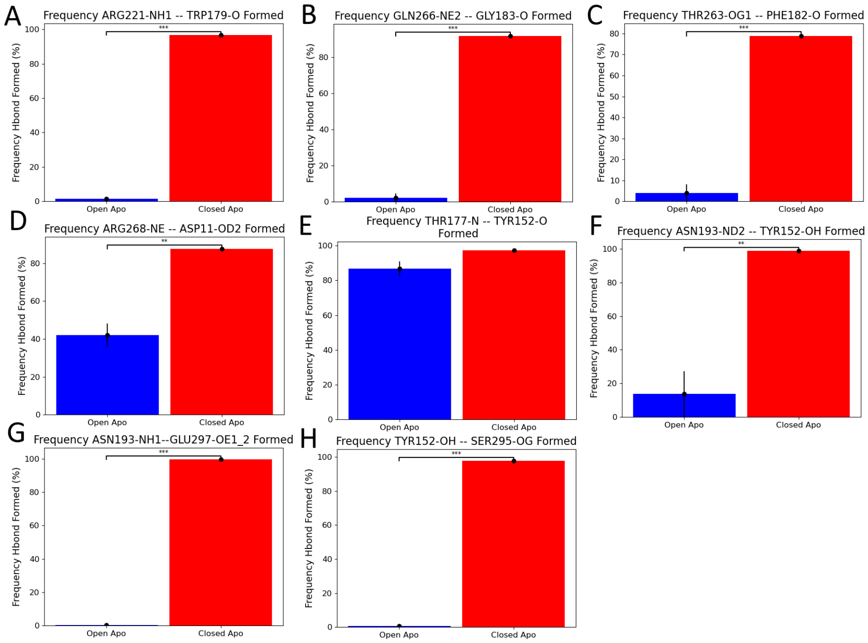


Figure S10: Hydrogen bonds mediate allosteric communication between the $\alpha 7$ helix and the active site. The apo open trajectories are initialized and maintain an open WPD loop with a disordered $\alpha 7$ helix, the apo closed trajectories are initialized and maintain a closed WPD loop with an ordered $\alpha 7$ helix. Bonds immediately interacting with the active site form between the P-loop and WPD-loop (A) and connecting the WPD-loop to the $\alpha 6$ helix (B–C). An additional bond forms between the $\alpha 6$ helix and the $\alpha 1$ helix (D). The bond between the the L11-loop and the WPD-loop is maintained regardless of $\alpha 7$ helix or WPD-loop conformation (E). Prior work indicates that this bond is important for mediating communication between the active and allosteric sites. The L-11 Loop is then connected to the $\alpha 3$ and $\alpha 7$ helices (F–H). (G) The bond between N193 and E297 could utilize either OE1 or OE2 on E297 as the electron acceptor. If a bond was formed between either electron acceptor it was counted towards the overall total. Statistically significant differences are denoted for $p < 0.01$ (**) and $p < 0.001$ (***). Error bars represent the standard error on the mean compared between trajectories with the given conformation (n=3 for apo open, n=3 for apo closed, and n=2 for b/w open+closed).

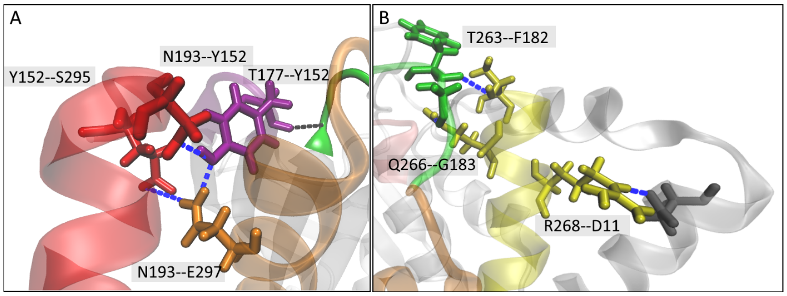


Figure S11: Our analysis identified differences in the H-bonds correlated to WPD loop conformation compared to previous studies [1, 4, 3]. (A) Near the allosteric site, previous studies identified the bonds between N193–Y152 and N193–E297 as those responsible for connecting the L-11 loop to the $\alpha7$ helix. Our study identified an additional bond Y152–S295 which directly connects the L-11 loop and the $\alpha7$ helix. (B) Previous networks have also identified T266–G183 which directly stabilizes the WPD loop through connection to the $\alpha6$ helix. This study identified two additional bonds which add an additional connection between the WPD-loop and the $\alpha6$ helix (T263–F182) and connecting the $\alpha6$ and $\alpha1$ helices (R268–D11). Flexibility and redundancy in H-bond networks is reasonably common in biological systems, though it may also reflect differences in procedures used for the simulation.

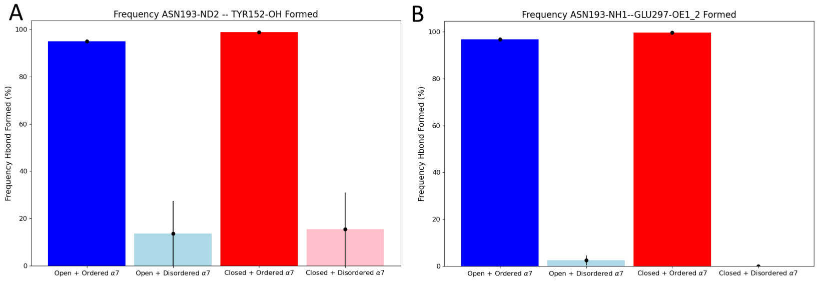


Figure S12: Of the h-bonds identified in the h-bond network only these two bonds were correlated specifically to the ordered structure of the $\alpha7$ helix. Even if the WPD loop was forced into a closed conformation with a disordered $\alpha7$ helix neither bond were able to form. This suggests that these h-bonds are the first within the network to be disrupted by the disordering of the $\alpha7$ helix upon ligand binding.

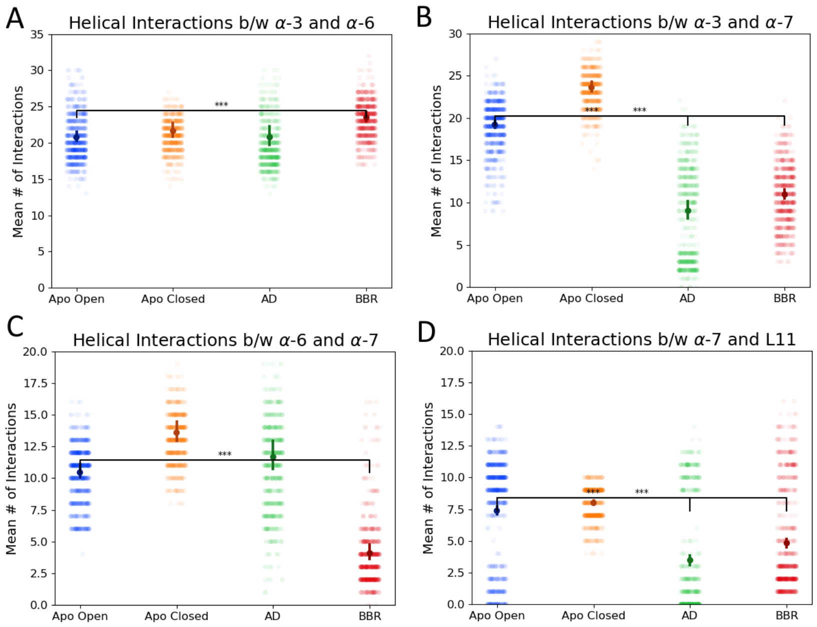


Figure S13: AD and BBR disrupt interactions within the helical triad that defines the allosteric site. (A) The interactions between the $\alpha3$ and $\alpha6$ helices increase by ~4% from the apo closed to apo open states. In the presence of AD, interactions between the $\alpha3$ and $\alpha6$ helices increased by ¡1% compared to the apo open state, and increase by 12% in the presence of BBR. In light of both (i) inconsistency in these changes between the two ligands and (ii) their small magnitude, they are likely not important to allosteric inhibition. (B) Interactions between the $\alpha3$ and $\alpha7$ helices decreased by 21% from the apo closed to open states, and decreased further by 72% and 55% for AD and BBR bound states, respectively. Both large magnitude of these changes and their consistency between inhibitors suggests that the disruption of these interactions is crucial to allosteric inhibition. (C) Interactions between the $\alpha6$ and $\alpha7$ helices increased by 26% from the apo closed to open states and increased further by 11% for AD while interactions decreased by and 87% for AD and BBR bound states, respectively. Due to the significant differences of the effect of the inhibitors on this interface, disruption of interactions at this interface is clearly not required for inhibition to occur. (D) Interactions between the $\alpha7$ helix and L-11 loop decreased by 8% from the apo closed to open states and decreased further by 71% with AD bound and 42% with BBR bound states. Given that these interactions are not significantly disrupted in the apo state it is possible this additional disruption is involved in the allosteric inhibition mechanism. The horizontal bars highlight statistical significance determined using Welch's T-test comparing the uncorrelated samples from trajectories fitting the given categories, $p \leq 0.001$.

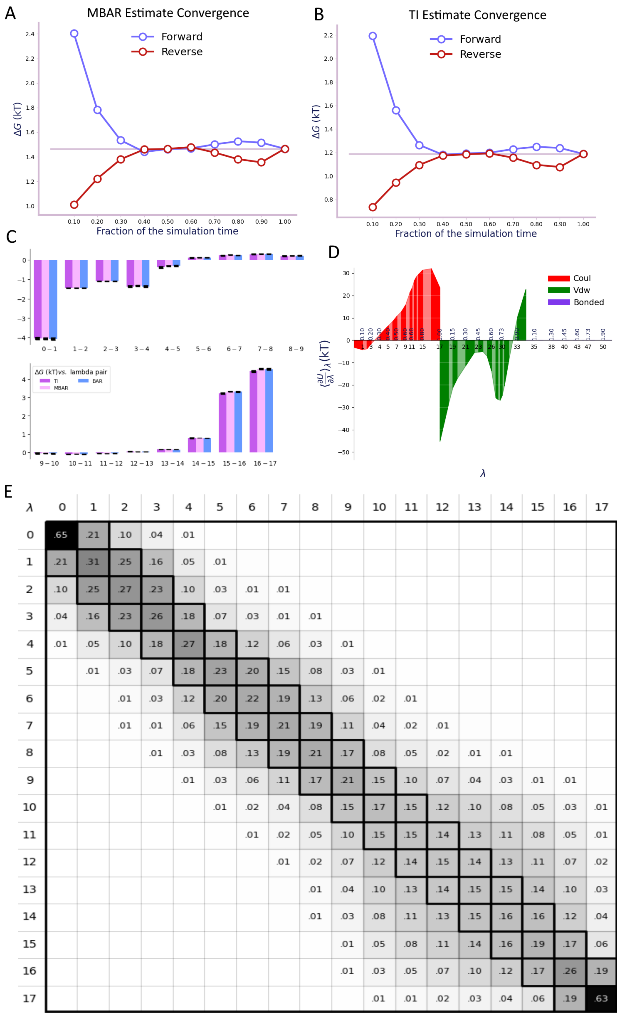


Figure S14: Optimization of relative free energy calculations for sufficient sampling and estimate convergence for the equilibrated trajectory. (A-B) We selected a simulation duration for which forward and reverse solvation free energy estimates for a test mutant (F196A) converged for both (A) MBAR and (B) TI estimates. Convergence was reached by comparing a minimum of the first and last 40% of the trajectory. This implies that the full equilibrated trajectory (the final 60% of this trajectory) is sampling the same free energy surface and is thus likely converged and the full equilibrated trajectory will be used for additional analysis. (C) We optimized the $\lambda$ state spacing to minimize error in ΔG between each pair of states and to ensure that TI, MBAR, and BAR estimators produced statistically indistinguishable results. (D) We verified that the lambda spacing produced a relatively smooth curve for $\frac{\partial U}{\partial \lambda}$ to avoid a high magnitude of $\frac{\partial^2 U}{\partial \lambda^2}$. This metric is particularly relevant for the accuracy of the TI estimate. (E) We verified overlap between adjacent $\lambda$ states to ensure that the sum of the off-diagonal values are $\geq 0.3$.

| Mutation | Forward Primer | Reverse primer |
|---|---|---|
| L192F | CAGCCTCATTCTTCAACTTTCTTTTCAAAGTC | GACTTTGAAAAGAAAGTTGAAGAATGAGGCTG |
| L195F | CATTCTTGAACTTTTTCTTCAAAGTCCGAGAG | CTCTCGGACTTTGAAGAAAAAGTTCAAGAATG |
| F196A | CTTGAACTTTCTTGCCAAAGTCCGAG | CTCGGACTTTGGCAAGAAAGTTCAAG |
| E276F | GCTGTGATCTTCGGTGCCAAATTC | GAATTTGGCACCGAAGATCACAGC |
| F280Y | CGAAGGTGCCAAATACATCATGGG | CCCATGATGTATTTGGCACCTTCG |
| V287T | GACTCTTCCACGCAGGATCAGT | ACTGATCCTGCGTGGAAGAGTC |

Table S1: Primers for Site-Directed Mutagenesis

| | WPD Conformation | $\alpha 7$ Helix | Ligand |
|---|---|---|---|
| 1 | Open | Absent | Apo |
| 2 | Closed | Absent | Apo |
| 3 | Open | Ordered | Apo |
| 4 | Closed | Ordered | Apo |
| 5 | Open | Disordered | Apo |
| 6 | Closed | Disordered | Apo |
| 7 | Open | Absent | Bound |
| 8 | Closed | Absent | Bound |
| 9 | Open | Ordered | Bound |
| 10 | Closed | Ordered | Bound |
| 11 | Open | Disordered | Bound |
| 12 | Closed | Disordered | Bound |

Table S2: The table describes the protein and ligand criteria used for MD simulations. We used all conformations of the WPD loop and $\alpha 7$ helix for AD, but only cases 9, 10, and 12 for BBR. All simulations carried out with the disordered $\alpha 7$ helix used four different conformations (See Materials and Methods).

Table S3: The accompanying Excel file (TableS3.xlsx) provides measurements of tryptophan fluorescence, including error and sample sizes, used for Figure 1B.

Table S4: The accompanying Excel file (TableS4.xlsx) provides initial rates of pNPP hydrolysis, including error and sample sizes, and estimates of the fractional change in inhibition (F) derived from these measurements. We used this data to construct Fig. 3B.

| $\alpha 7$ Initialized | WPD Initialized | Loc1 | Loc2 | Loc3 | Loc4 | % Unbound | Other Bound |
|---|---|---|---|---|---|---|---|
| ordered | open | 14% | 0% | 66% | 22% | 0% | 12% |
| ordered | closed | 0% | 0% | 59% | 40% | 0% | 1% |
| absent | open | 0% | 0% | 34% | 63% | 1% | 2% |
| absent | closed | 0% | 2% | 0% | 85% | 0% | 13% |
| disordered 1 | open | 1% | 1% | 1% | 1% | 84% | 12% |
| disordered 1 | closed | 0% | 0% | 0% | 1% | 60% | 40% |
| disordered 2 | open | 0% | 0% | 74% | 1% | 11% | 14% |
| disordered 2 | closed | 0% | 3% | 6% | 0% | 61% | 30% |
| disordered 3 | open | 67% | 18% | 0% | 0% | 0% | 15% |
| disordered 3 | closed | 20% | 80% | 0% | 0% | 0% | 0% |
| disordered 3 | closed | 1% | 3% | 1% | 0% | 92% | 3% |
| disordered 4 | closed | 99% | 1% | 0% | 0% | 0% | 0% |
| disordered 4 | closed | 100% | 0% | 0% | 0% | 0% | 0% |

Table S5: The binding conformation of AD in each equilibrated trajectory. All percentages are rounded to the nearest whole number. Loc1 corresponds to the crystal binding location. Alternative binding locations are defined in the Methods and Materials section and shown in Figure 3. The "other bound" category is for when the ligand is not bound to one of the defined binding location but it still interacting with the protein. These binding locations are fleeting and inconsistent between trajectories. For the disordered 3 and 4 $\alpha 7$ helix structures two replicas were completed with the same initial configuration with a closed WPD loop but distinct random seed. For a full description of the determination of occupancy percentages see section S1.6.

| | TI Solvation FE | TI Complex FE | TI Binding FE | MBAR Solvation FE | MBAR Complex FE | MBAR Binding FE |
|---|---|---|---|---|---|---|
| F196A | 0.48 ± 0.066 | 0.917 ± 0.067 | 0.437 ± 0.133 | 0.742 ± 0.055 | 1.123 ± 0.054 | 0.381 ± 0.109 |
| L192F | 25.165 ± 0.052 | 24.79 ± 0.051 | -0.375 ± 0.103 | 25.031 ± 0.047 | 24.734 ± 0.048 | -0.297 ± 0.095 |
| L195F | 25.922 ± 0.043 | 25.878 ± 0.043 | -0.044 ± 0.086 | 25.959 ± 0.037 | 25.94 ± 0.037 | -0.019 ± 0.074 |
| F280Y | -25.082 ± 0.066 | -24.448 ± 0.068 | 0.634 ± 0.134 | -22.317 ± 0.039 | -22.081 ± 0.033 | -0.236 ± 0.072 |
| E276F | 93.634 ± 0.167 | 92.108 ± 0.163 | -1.526 ± 0.33 | 78.032 ± 0.386 | 77.659 ± 0.308 | 0.373 ± 0.694 |
| V287T | -14.581 ± 0.11 | -14.525 ± 0.076 | 0.056 ± 0.186 | -11.05 ± 0.063 | -11.076 ± 0.043 | -0.026 ± 0.106 |

Table S6: This table depicts the FE estimates from the TI and MBAR estimators and their corresponding errors as produced by alchemlyb analysis (Error description in Methods). The binding FE estimate is the difference between the complex FE estimate and the solvation FE estimate.
